# Proteostasis as a fundamental principle of Tau immunotherapy

**DOI:** 10.1101/2024.02.12.580007

**Authors:** Esteban Cruz, Rebecca M. Nisbet, Pranesh Padmanabhan, Ashley J. van Waardenberg, Mark E. Graham, Godfrey Nkajja, Swara Tapaswi, Phil Robinson, Jürgen Götz

**Author notes:** Correspondence to: J.G., Clem Jones Centre for Ageing Dementia Research (CJCADR), Queensland Brain Institute (QBI), The University of Queensland, St Lucia Campus (Brisbane), QLD 4072, Australia.

## Abstract

The microtubule-associated protein Tau is a driver of neuronal dysfunction in Alzheimer’s disease and numerous other tauopathies. In this process, Tau initially undergoes subtle changes to its abundance, subcellular localisation and a vast array of post-translational modifications including phosphorylation, that progressively result in the protein’s aggregation and dysregulation of multiple Tau-dependent cellular processes.

Given the various loss- and gain-of-functions of Tau in disease and the brain-wide changes in the proteome that characterise tauopathies, we asked whether targeting Tau would restore the alterations in proteostasis observed in disease.

To this end, we generated a novel pan-Tau antibody, RNJ1, that preferentially binds human Tau and neutralises proteopathic seeding activity in multiple cell lines and benchmarked it against a clinically tested pan-Tau antibody, HJ8.5 (murine version of tilavonemab). We next evaluated both antibodies, alone and in combination, in the K3 mouse model of tauopathy, showing reduced Tau pathology and improvements in neuronal function following 14 weekly treatments, without obtaining synergistic effects for the combination treatment.

To gain insight into molecular mechanisms contributing to improvements in neuronal function, we employed quantitative proteomics and phosphoproteomics to first establish alterations in K3 mice relative to WT controls at the proteome level. This revealed 342 proteins with differential abundance in K3 mice, which are predominantly involved in metabolic and microtubule-associated processes, strengthening previously reported findings of defects in several functional domains in multiple tauopathy models. We next asked whether antibody-mediated Tau target engagement indirectly affects levels of deregulated proteins in the K3 model. Importantly, both immunotherapies, in particular RNJ1, induced abundance shifts in this protein subset towards a restoration to wild-type levels (proteostasis). A total of 257 of 342 (∼75.1%) proteins altered in K3 were closer in abundance to WT levels after RNJ1 treatment. The same analysis indicated a similar response in K3 mice treated with HJ8.5, with approximately 72.5% of these altered proteins also showing changes in the same direction as wild-type. Furthermore, analysis of the phosphoproteome showed an even stronger restoration effect with RNJ1, with ∼82.1% of altered phosphopeptides in K3 showing a shift to WT levels, and 75.4% with HJ8.5. Gene set over-representation analysis (ORA) further confirmed that proteins undergoing restoration are involved in biological pathways affected in K3 mice. Together, our study suggests that a Tau immunotherapy-induced restoration of proteostasis links target engagement and treatment efficacy.

## Introduction

A histopathological hallmark not only of Alzheimer’s disease (AD) but also of more than two dozen neurodegenerative diseases, collectively termed tauopathies, is the intracellular aggregation of Tau, a neuronally enriched protein. AD is further characterised by a second hallmark lesion in the form of amyloid-β (AΒ)-containing plaques in the extracellular space,^1^ with both pathologies impairing multiple cellular functions.^2–6^ In the pursuit of clearing these aggregates, substantial efforts have been invested into developing active and passive immunotherapies targeting the two molecules.^7,8^ This has resulted in the recent approval of two anti-AΒ antibodies, aducanumab and lecanemab, by the US Food and Drug Administration (FDA), with additional antibodies following suit.^9–11^ In contrast, Tau antibodies have, to date, failed to demonstrate clinical efficacy, necessitating the development of more potent Tau-targeting antibodies.

Developing an effective Tau immunotherapy, however, poses significant challenges. Tau is highly heterogeneous as in the human brain the protein exists as six isoforms that undergo a myriad of post-translational modifications, including hyperphosphorylation, and how the different forms of Tau each contribute to pathogenicity across tauopathies is not fully understood.^1^ Furthermore, Tau is differentially distributed across distinct neuronal sub-compartments and is also found in the extracellular milieu and glial cells. Adding to this complexity, Tau pathology varies significantly between the transgenic models that are used as validation tools for therapeutic interventions.^12^ This makes it challenging if one explores an anti-Tau antibody in a transgenic mouse model and focuses the validation largely or solely on changes to abundance and specific phosphorylation of Tau, rather than on the impairments that are elicited at multiple and often subtle levels in different functional domains.

In this study, we generated a novel Tau antibody, RNJ1, and compared it in vitro and in vivo with the clinically tested Tau antibody HJ8.5 (tilavonemab).^13^ We first validated the antibodies’ capacity to bind monomeric and aggregated forms of human Tau and to neutralise aggregation induced by Tau seeds from tauopathy mice and human AD tissue in two independent Tau biosensor cell lines.^14^ We next found that RNJ1, but not HJ8.5, reduced total Tau levels and improved behavioural deficits in the K369I mutant human Tau transgenic model K3. This strain displays Tau pathology and pronounced motor impairment with an early age-of-onset at 4 weeks, that manifests as a lack of locomotor ability and coordination that can be readily assessed by the Rotarod test. Given that protein changes in AD and animal models are proxies for neuronal dysfunction,^6,15,16^ in addition to focusing solely on changes to tau pathology, we conducted proteomic and phospho-proteomic analyses to investigate how the Tau antibodies induce multiple subtle changes to Tau and its interactions and thereby collectively contribute to improvements in different functional domains. Importantly, we found that RNJ1 shifts the balance in multiple domains towards reinstalling a wild-type state of homeostasis in K3 mice. This shift elicited by RNJ1 treatment was induced at the level of both the proteome and the phospho-proteome. Our study highlights the potential for using proteomic changes as a correlate of Tau immunotherapeutic efficacy and advocates RNJ1 for clinical testing.

## Materials and methods

The following materials and methods are described in the Supplement: Antibodies

Preparation of recombinant human Tau and single chain variable fragment (scFv) library panning

Preparation of monoclonal scFvs for ELISA

Purification of scFvs and western blot analysis

Generation of mouse IgG RNJ1

Epitope mapping of RNJ1 and confirmation of HJ8.5 epitope through indirect ELISA

Determination of antibody EC_50_ against monomeric hTau-441

Determination of antibody binding curves against aggregated sarkosyl-insoluble Tau

Surface plasmon resonance

Generation of Tau RD P301S SH-SY5Y FRET biosensor cells

Cell culture and Tau seed neutralisation assays

Large scale production of RNJ1 and HJ8.5 for treatment study

Animals, immunisation and behavioural tests

Tissue processing for immunofluorescence

Immunofluorescence imaging and image analysis

Protein extraction from whole brain lysates and western blot analysis

Sample preparation for mass spectrometry

LC-MS/MS analysis

MS data processing

MaxQuant data processing of the proteome and phosphoproteome of the antibody treatment study

Protein kinase activity prediction

Other bioinformatic and statistical analysis

## Results

### Isolation of the Tau antibody RNJ1 from a human phage display library and epitope mapping

To isolate a Tau-specific antibody, the Tomlinson I + J human synthetic single-fold single-chain variable fragment (scFv) library was panned in multiple rounds against full-length human recombinant Tau (hTau-441), yielding an enrichment of positive scFvs including RNJ1 (Supplementary Fig. 1A-D). This scFv bound to recombinant hTau-411 (the longest human brain Tau isoform), by ELISA and by western blot analysis (Supplementary Fig. 1E,F). To further characterise RNJ1 and determine its preclinical efficacy, its variable heavy and variable light chains were reformatted into a mouse IgG1 backbone containing a kappa light chain.

Using sequentially truncated forms of recombinant hTau-441 (Fig. 1A), we mapped the RNJ1 epitope by indirect ELISA, and also confirmed the epitope of the clinically tested Tau antibody HJ8.5,^13^ which we cloned and purified for benchmarking purposes. Both RNJ1 and HJ8.5 bound exclusively to forms of Tau containing the first 44 amino acids (aa); however, whereas RNJ1 had a binding site within aa 1-22, HJ8.5 recognised the variant containing aa 23-44, consistent with its reported epitope (aa 25-30) (Fig. 1B,C).^13^ Next, we employed an overlapping synthetic peptide library spanning aa 1-22 with peptide lengths of 10 aa and an offset of 2 aa to map the RNJ1 epitope at higher resolution (Fig. 1D). The peptides were conjugated to bovine serum albumin (BSA) and used as antigens in an indirect ELISA. Maximal binding was detected for peptides spanning aa 9-18 and 11-20, with residual binding observed to aa 13-22. Similarly, surface plasmon resonance (SPR) using the BSA-conjugated peptides as ligands and RNJ1 as analyte showed maximal binding for aa 9-18 and 11-20, with some binding to aa 13-22, and nearly no detectable signal for the remaining peptides (Supplementary Fig. 2). Together, these results show that RNJ1 binds to a segment of Tau spanning aa 9-22.

**Figure 1.**
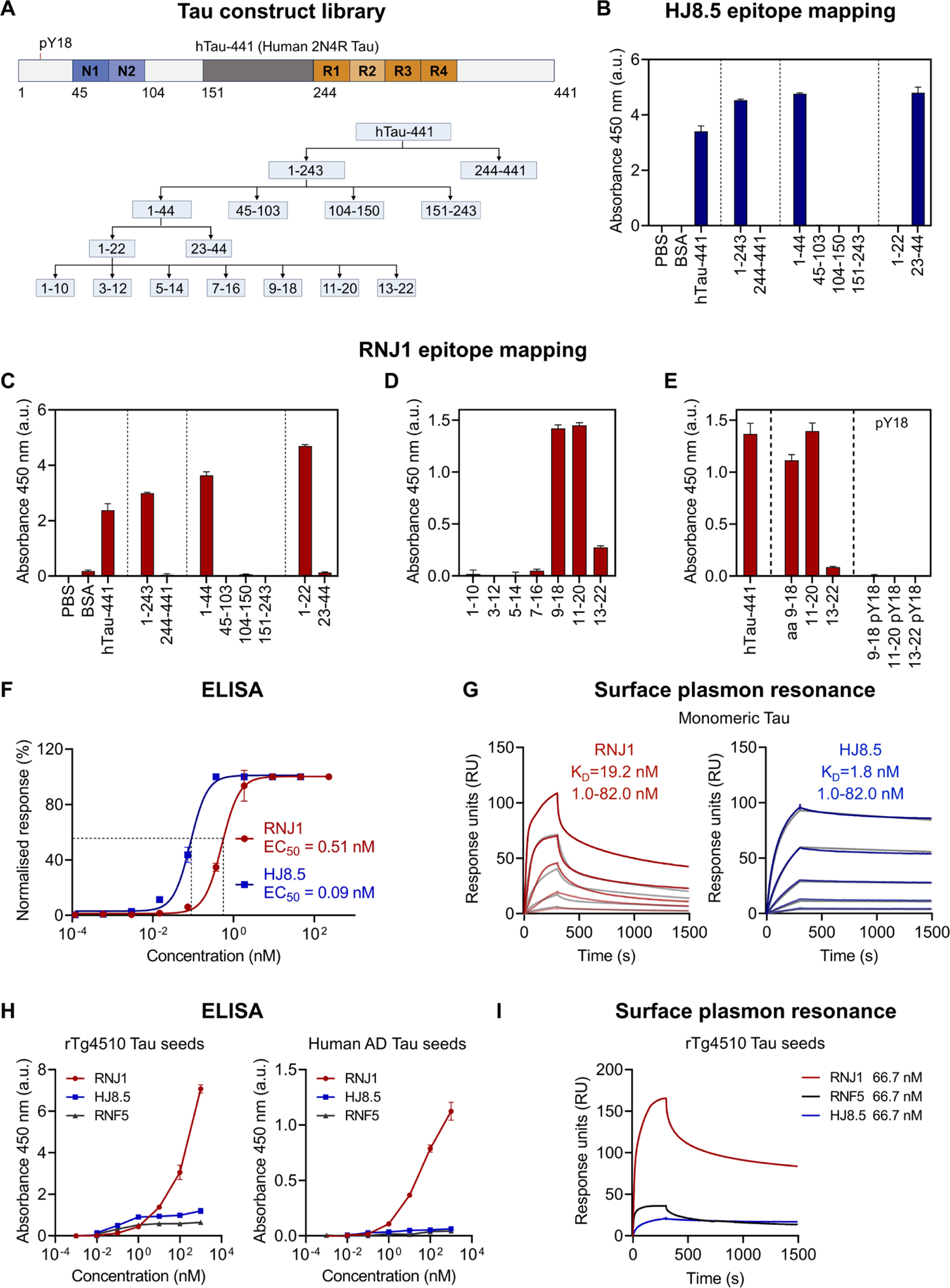
Antibody RNJ1 binds to an amino-terminal epitope of Tau and has stronger reactivity to aggregated Tau than the amino-terminal antibodies, HJ8.5 and RNF5. (**A**) Schematic depicting Tau fragments expressed as fusion proteins with maltose-binding protein for epitope mapping of novel antibody RNJ1 and benchmarking antibody HJ8.5. (**B**) Epitope mapping of HJ8.5 by indirect ELISA using the Tau fusion proteins as antigens. (**C**) Epitope mapping of RNJ1 by indirect ELISA with the Tau fusion proteins. (**D**) Higher resolution mapping using 10 amino acid (aa) long overlapping peptides of Tau’s amino-terminal 22 aa conjugated to BSA (middle panel). (**E**) Indirect ELISA epitope mapping using BSA-conjugated non-phosphorylated and phosphorylated peptides (right panel). (**F**) Determination of RNJ1 and HJ8.5 concentration yielding half-maximal binding, EC_50_, to monomeric recombinant human 2N4R Tau (hTau-441) by indirect ELISA. (**G**) Surface plasmon resonance (SPR) sensorgrams of RNJ1 and HJ8.5 (analytes) binding to monomeric recombinant hTau-441. Sensorgrams shown for antibody injections at 1.0, 3.0, 9.1, 27.3 and 82.0 nM. Both sensorgrams were fitted to a bivalent binding model. Experimental curves for RNJ1 are shown in red and for HJ8.5 in blue, and model fits in gray. (**H**) Binding curves of RNJ1, HJ8.5 and RNF5 from indirect ELISAs using sarkosyl-insoluble rTg4510 transgenic and human AD Tau seeds as antigens. (**I**) SPR sensorgrams of RNJ1, HJ8.5 and RNF5 (analytes) binding to sarkosyl-insoluble Tau seeds derived from rTg4510 mice. In B-C, and E, n = 4 technical replicates. In D, F and H, n = 3 technical replicates. Data shown as mean ± SEM.

In addition, given that tyrosine 18 (Y18) is a substrate of Fyn and phosphorylation of this epitope is found in tauopathy and in animal models of tauopathy,^17,18^ we tested RNJ1 reactivity against the synthetic peptides aa 9-18, 11-20 and 13-22 phosphorylated at Y18. However, we did not detect binding, suggesting that RNJ1 binds to a segment spanning aa 9-22 that is not phosphorylated at Y18 (Fig. 1E).

### RNJ1 displays stronger reactivity to aggregated Tau than HJ8.5 and RNF5

Given that our epitope mapping revealed adjacent epitopes for RNJ1 (aa 9-22) and HJ8.5 (aa 25-30) within Tau’s amino-terminal domain, we next interrogated whether this difference affects the binding profile to monomeric and aggregated Tau. We first estimated the half-maximal binding concentration, EC_50_, through indirect ELISAs using recombinant hTau-441 in a monomeric form as antigen, and found that the EC_50_ of HJ8.5 (0.09 nM, 95% CI: 0.077-0.093 nM) was approximately five times lower than that of RNJ1 (0.51 nM, 95% CI: 0.46-0.57 nM) (Fig. 1F). We then measured the binding kinetic constants, using monomeric hTau-441 as ligand and the two antibodies at various concentrations as analytes in multi-cycle kinetic SPR analyses. Based on SPR sensorgrams, the antibodies displayed a good fit with a bivalent analyte model, with HJ8.5 showing higher affinity (equilibrium dissociation constant, *K_D_* = 1.8 nM) to monomeric Tau than RNJ1 (*K_D_* = 19.15 nM) (Fig. 1G).

To assess binding to aggregated Tau, we coated ELISA plates with sarkosyl-insoluble Tau obtained from P301L mutant Tau transgenic rTg4510 mice^19^ and human AD brain lysates, and tested, in addition to RNJ1 and HJ8.5, a third amino-terminal antibody we have described previously, RNF5, which exhibits specificity for aa 35-44.^20^ HJ8.5 and RNF5 both reached binding saturation at low antibody concentrations, whereas RNJ1 did not reach saturation within the concentration range tested, resulting in a significantly higher maximal response against sarkosyl-insoluble Tau from both mouse and human brain (Fig. 1H). To gain further insight into the binding properties of the antibodies to aggregated Tau, we performed SPR using sarkosyl-insoluble rTg4510 Tau as ligand and the antibodies as analytes. While the structural complexity of aggregated Tau precludes deriving binding kinetic constants with high confidence in SPR, it was evident from the sensorgrams that RNJ1 produced a much higher binding signal at equivalent antibody concentrations than HJ8.5 and RNF5 (Fig. 1I), reflecting the higher reactivity of RNJ1 against aggregated Tau determined by indirect ELISA (Fig. 1H). Together, the differential binding profile of RNJ1 to aggregated Tau determined by ELISA and SPR suggested that RNJ1 could have a higher capacity to neutralise templated aggregation induced by proteopathic Tau seeds.

### RNJ1 shows remarkable efficiency at neutralising proteopathic Tau seeds *in vitro*

Given that in AD, pathological Tau has seeding capacity,^21^ and that RNJ1 displayed increased reactivity against aggregated Tau compared with HJ8.5 and RNF5, we sought to determine whether RNJ1 would have a higher capacity to neutralise Tau seeds in a biosensor system. For this, we employed the widely used Tau RD P301S FRET HEK293 (human embryonic kidney) cells which co-express the microtubule-binding repeat domain (RD) of P301S mutant Tau, fused to cyan fluorescent protein (CFP) or yellow fluorescent protein (YFP). In this system, internalisation of exogenous Tau seeds induces aggregation of the recombinant RD Tau expressed by the cells, producing a Förster resonance energy transfer (FRET) signal that can be measured by microscopy or flow cytometry.^22^

We treated the HEK293 biosensor cells with sarkosyl-insoluble rTg4510 Tau seeds in the presence or absence of the antibodies to assess the effect of antibody treatment on the formation of Tau inclusions, and measured the FRET signal by flow cytometry (Fig. 2A,B). Using Tau seeds at 200 ng/mL, we tested the antibodies at concentrations from 10 to 300 nM. When comparing the effects of the antibodies on Tau seeded aggregation, RNJ1 treatment was most effective at decreasing the mean integrated fluorescence intensities (IFI) for the FRET signal across all concentrations tested, reducing the IFI to ∼30% of the ‘no antibody’ control at the highest antibody concentration of 300 nM, compared with ∼42% for HJ8.5 and ∼70% for RNF5 (Fig. 2A-C).

**Figure 2.**
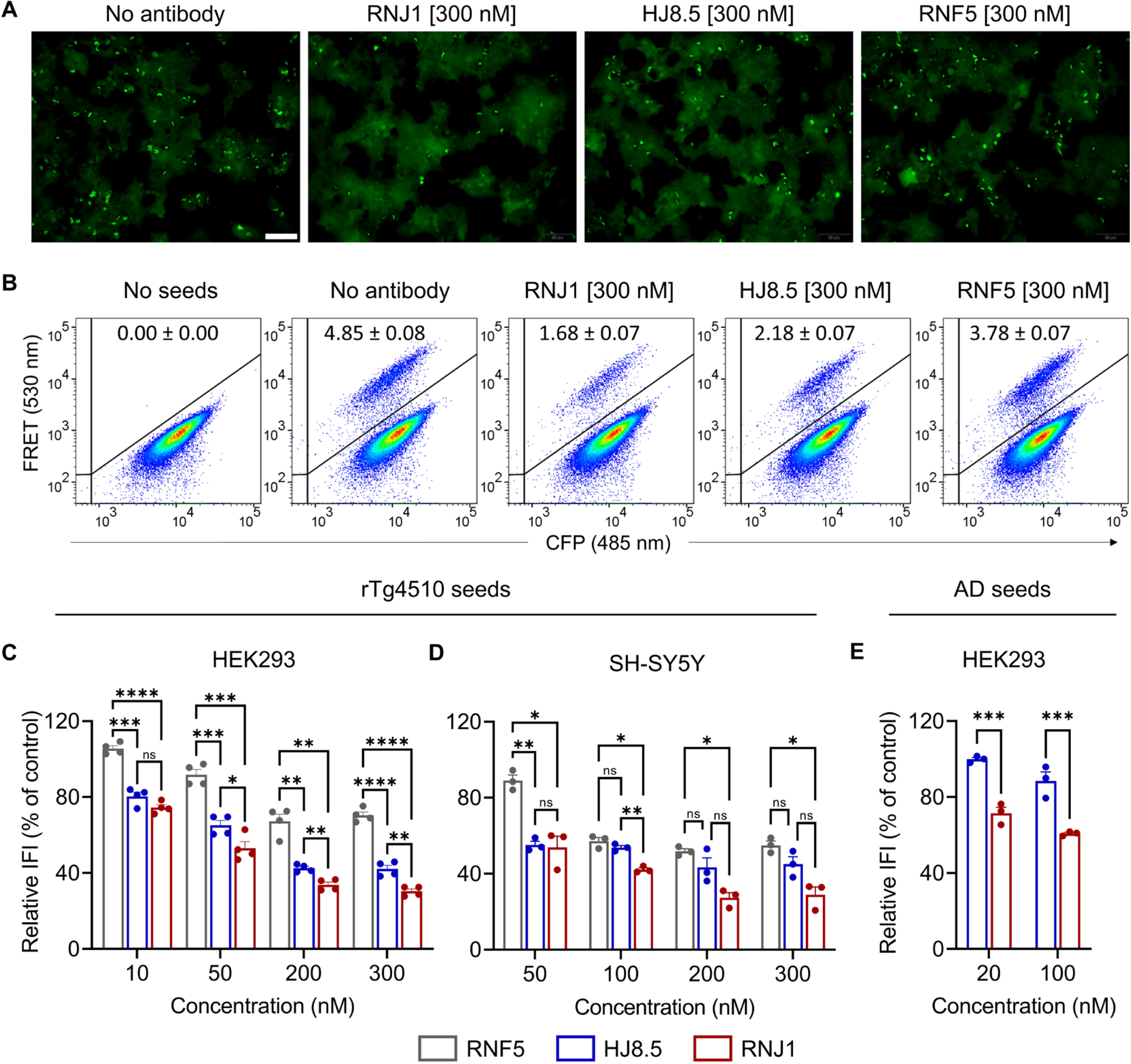
RNJ1 inhibits seeding activity of sarkosyl-insoluble Tau in two distinct Tau biosensor cells. (**A**) Representative epifluorescence microscopy images of Tau RD P301S FRET HEK293 biosensor cells 48 h post-exposure to rTg4510 transgenic mouse-derived Tau seeds in the presence or absence of 300 nM RNJ1, HJ8.5 or RNF5. All samples were treated with the same amount of Tau seeds. (**B**) Representative FRET-based flow cytometry scatter plots of the HEK293 biosensor cells treated with rTg4510 Tau seeds with and without antibody. The percentage of FRET-positive cells is shown as mean ± SEM. No FRET signal was detected in biosensor cells without Tau seed exposure. Scale bar represents 100 µm. (**C**) HEK293 biosensor cells treated with rTg4510 seeds and Tau antibodies. The percentage reduction in integrated FRET intensity values from flow cytometry analysis relative to the no antibody control values is shown. (**D**) Complementary Tau seed neutralization assay performed in a novel Tau RD P301S SH-SY5Y FRET biosensor system. (**E**) Seeding neutralization assay in HEK293 cells performed as in C but with AD Tau seeds. In B and C, n = 4 technical replicates for each concentration, 50,000 cells analyzed per replicate. In D and E, n = 3 technical replicates for each concentration, 50,000 cells analyzed per replicate. Statistical analysis was performed using two-way ANOVA with the Holm-Sidak’s multiple comparison correction. *p < 0.05, **p < 0.01, ***p < 0.001, ****p < 0.0001.

We next used a complementary FRET-biosensor system we had established in human SH-SY5Y neuroblastoma cells to assess neutralisation of Tau seeding in a more neuron-like model. Again, RNJ1 displayed superior neutralisation capacity, decreasing the IFI of the FRET signal to ∼29% of the ‘no antibody’ control at the highest antibody concentration tested (300 nM) compared with ∼45% and ∼55% for HJ8.5 and RNF5, respectively (Fig. 2D). The mean IFI was consistently lower for RNJ1 than for HJ8.5 across all concentrations tested, although the difference achieved statistical significance only at 100 nM. Of note, RNJ1 was significantly superior to RNF5 across the four tested concentrations.

We then asked whether the antibodies were also capable of neutralising AD brain-derived Tau seeds. We therefore treated the HEK293 biosensors with sarkosyl-insoluble AD Tau (Fig. 2E). Treatment with 100 nM RNJ1 reduced seeding intensity to nearly ∼60% of the seeding detected with the ‘no antibody’ control, whereas inhibition with equivalent doses of HJ8.5 only achieved a reduction to ∼88%. Treatment with RNJ1 achieved significantly higher seeding inhibition relative to HJ8.5 at both 20 and 100 nM concentrations. Together, these results show that RNJ1 exhibits a superior capacity to neutralise Tau seeds obtained from both a tauopathy mouse model and human AD brain tissue. Importantly, because the three tested antibodies bind to an amino-terminal epitope which is lacking in the Tau RD constructs expressed in the biosensor cells, the observed seeding inhibition is likely to occur by binding to the exogenous Tau seeds rather than to the RD Tau expressed by the biosensor cells.

### RNJ1 treatment improves motor functions in the K3 mouse model of tauopathy

Having established that HJ8.5 and RNJ1 differ in their binding profiles to and neutralisation capacity of seed-competent Tau, we next compared their efficacy, alone and together, in a longitudinal study using the K369I frontotemporal dementia-mutant Tau transgenic K3 mouse model of tauopathy.^23^ K3 mice display Tau pathology and pronounced motor impairment with an early age-of-onset at 4 weeks, that manifests as a lack of locomotor ability and coordination as assessed, for example in the Rotarod test.^24^

For treatment group allocation, forty-eight K3 mice and twelve age-matched wild-type (WT) littermates (even male-to-female ratio) were subjected to a Rotarod baseline test at 4 weeks of age, and then allocated based on an equivalent average performance to five treatment groups (Fig. 3A): K3 mice treated with RNJ1 (K3^RNJ1^), HJ8.5 (K3^HJ8.5^), an antibody combination (K3^R+H^) and vehicle (K3^Veh^), plus vehicle-treated WT mice (WT^Veh^). The mice received 14 weekly intraperitoneal doses of the antibody at 50 mg/kg for the single treatments and 25 mg/kg each for the combination, starting at 5 weeks of age. Rotarod performance was tracked longitudinally every 4 weeks, with a final test at the end of treatment. The K3 mice in all groups showed a significantly impaired performance in the Rotarod test compared to WT^Veh^, with WT^Veh^ mice reaching a maximum Latency-to-fall (180 s) in nearly every test session (Fig. 3B, top). K3^RNJ1^ mice displayed a higher mean Latency-to-fall than all other K3 groups at every test session after treatment start, but these differences did not reach statistical significance (Fig. 3B, top). To assess the overall performance of the groups over the course of treatment, we also calculated the area under the curve (AUC) of the Latency-to-fall. Notably, K3^RNJ1^ mice showed a 27 percent increase in the mean AUC compared to K3^Veh^, without reaching statistical significance (*P* = 0.0911) (Fig. 3B, bottom). K3^HJ8.5^ and K3^R+H^ mice showed no improvements throughout treatment or at time of completion compared to K3^Veh^ (Fig. 3B). Interestingly, we found that K3^Veh^ females performed better than K3^Veh^ males, with the second and third Rotarod tests (8 and 12 weeks of age) displaying significant differences (Fig. 3C, top), and the AUC being significantly higher for K3^Veh^ females (Fig. 3C, bottom). This prompted us to stratify the treatment groups based on sex to increase the sensitivity of the behavioural read-out (Fig. 3D).

**Figure 3.**
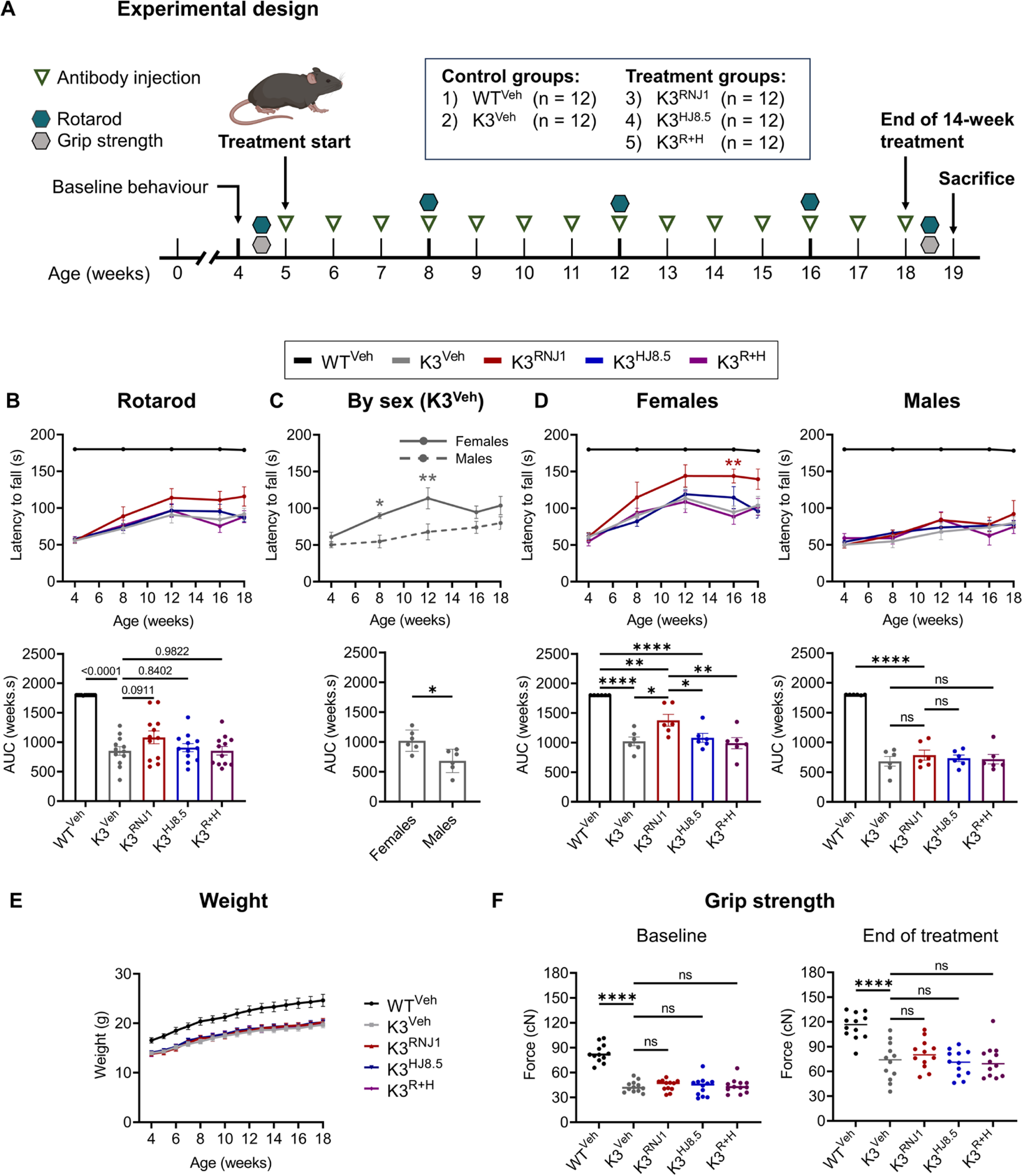
RNJ1 treatment improves motor function in female K3 mice. (**A**) Experimental design of longitudinal treatment study: Baseline behaviour and locomotor function were determined in 4 week-old K369I Tau mutant K3 mice with a Tau pathology and WT littermate controls to assign mice into experimental groups with equivalent average Rotarod performance. The experimental groups were vehicle-treated WT mice (WT^Veh^), and K3 mice treated with vehicle (K3^Veh^), RNJ1 (K3^RNJ1^), HJ8.5 (K3^HJ8.5^) or an antibody combination (K3^R+H^). Treatments were done weekly, and Rotarod performance was assessed every 4 weeks for 14 weeks. (**B**) Rotarod mean Latency-to-fall (top panel) for all study arms across the treatment period (*n* = 12 mice per group) were compared by calculating the corresponding area under the curve (AUC) (bottom panel). (**C**) Mean Latency-to-fall off the Rotarod displayed separately for male and female mice in the K3^Veh^ group (top panel), and corresponding AUC (bottom panel). Female K3 mice showed improved locomotor function and balance compared to K3 males (*n* = 6 mice per group). (**D**) Group-based sex-stratified Mean Latency-to-fall across the treatment period (*n* = 6 mice per group) (top panel) and corresponding AUC (bottom panel). (E-G). Female K3^RNJ1^ mice showed improved locomotor performance compared with female K3^Veh^ mice. (**E**) Mean weight per group across the study. (**F**) Mean forelimb grip strength per group at baseline and at the end of treatment. In (B-F) data are represented as mean ± SEM. Statistical comparisons for Rotarod AUC and grip strength in the treatment groups were performed using one-way ANOVA with Holm-Sidak’s multiple comparison correction, or using an unpaired t-test for comparison of Rotarod AUC based on sex in C. Comparison of the Rotarod Latency-to-fall across timepoints was performed with a two-way ANOVA with Holm-Sidak’s multiple comparison correction. \**P* < 0.05, \*\**P* < 0.01, \*\*\**P* < 0.001, \*\*\*\**P* < 0.0001.

Sex-stratified analysis of the treatment groups revealed that K3^RNJ1^ females significantly improved in their Rotarod performance, with the second and third tests (8 and 12 weeks of age) showing a trend towards improvement, that reached statistical significance by the fourth test (16 weeks of age) (Fig. 3D, top left). This improvement was also reflected by a statistically significant increase in the AUC of Latency-to-fall (Fig. 3D, bottom left). In contrast, the treated males showed no improvements (Fig. 3D, top right, and bottom right). Weight (Fig. 3E) and grip strength averages (Fig. 3F) were significantly different between WT and all K3 groups, with no treatment effect. Together, our findings demonstrate that RNJ1 effectively improved motor function in female K3 mice.

### RNJ1 treatment causes subtle reductions in total Tau levels in brain homogenates of female K3 mice

Given that the RNJ1 treatment improved Rotarod performance in female K3 mice, we evaluated the biochemical and histopathological changes induced by the antibody treatment in the female cohort. Tau accumulates and undergoes hyperphosphorylation in disease.^25^ We therefore assessed immunoreactivity for total (mouse and human) Tau and for Tau phosphorylated at the pathological AT8 epitope (pS202/pT205) by western blot analysis using whole brain lysates (Fig. 4A). This revealed a subtle, yet statistically significant mean reduction of ∼12% in total Tau in the K3^RNJ1^ group compared with K3^Veh^ (*P* = 0.0102) (Fig. 4A,B), with the K3^R+H^ group also showing a statistically significant reduction of 8.5% in total Tau (*P* = 0.049). Levels of AT8 displayed greater variability, with K3^RNJ1^ mice showing a ∼27% reduction; however, this reduction did not reach statistical significance (*P* = 0.1069) (Fig. 4A,B).

**Figure 4.**
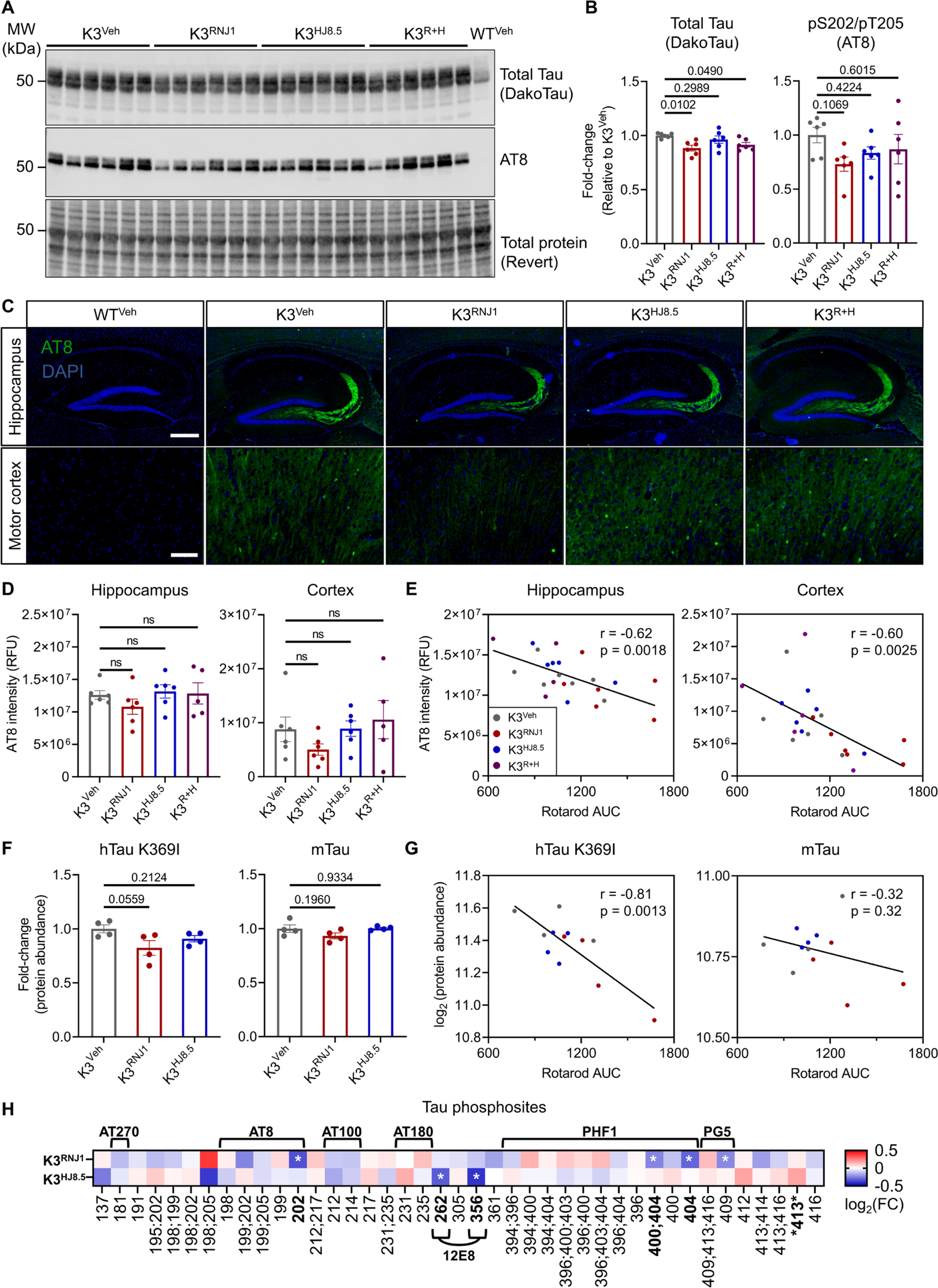
Total and phospho-tau levels are reduced in RNJ1-treated K3 mice. (**A**) Immunoblots of whole-brain lysates from female K3 mice probed for total Tau (DakoTau) and S202/T205-phosphorylated Tau (AT8). (**B**) Quantification of A using total protein stain (Revert^TM^ 700) for normalisation, represented as fold-change relative to the K3^Veh^ control group (*n* = 6 per group). (**C**) Representative (20x) images of AT8 immunoreactivity (green) with DAPI counterstaining for nuclei (blue) in the hippocampus and motor cortex of female mice. Scale bars represent 400 µm (top) and 100 µm (bottom). (**D**) Quantification of integrated fluorescence intensity in hippocampus and cortex in C. K3^Veh^, *n* = 6; K3^RNJ1^, *n* = 6; K3^HJ8.5^, n = 6; K3^R+H^, *n* = 5 (1 mouse excluded due to tissue damage). (**E**) Rotarod performance across the treatment study (Rotarod AUC) reveals a moderate inverse correlation with AT8 immunoreactivity for the hippocampus (left panel) (Pearson’s correlation *r* = -0.62, *P* = 0.0018) and cortex (right panel) (Pearson’s correlation *r* = -0.60, *P* = 0.0025). (**F**) Mass spectrometry quantification of protein abundance for the K369I hTau transgene and endogenous mouse Tau (*n* = 4 female mice per group). (**G**) Levels of hTau K369I show a strong inverse correlation with performance in Rotarod test (Pearson’s correlation *r* = -0.81, *P* = 0.0013), whereas endogenous mouse Tau levels only show a weak inverse correlation (Pearson’s correlation *r* = -0.32, *P* = 0.32). (**L**) Heatmap of Tau phosphopeptides detected through quantitative mass spectrometry. Phosphopeptides with fold-changes that achieved *P* < 0.05 are highlighted with an asterisk. Data in the heatmap represented as mean log_2_ (fold-change) relative to K3^Veh^. All other data represented as mean ± SEM. Statistical comparisons were performed with one-way ANOVA with Holm-Sidak’s post hoc multiple comparison’s test. \**P* < 0.05, \*\**P* < 0.01, \*\*\**P* < 0.001, \*\*\*\**P* < 0.0001.

We then assessed AT8 reactivity by immunofluorescence in motor cortex and hippocampus (Fig. 4C), brain areas characterised by Tau deposition in K3 mice.^20,24^ The mossy fibers of dentate granule cells, which innervate hilar cells and CA3 pyramidal cells, exhibit particularly pronounced reactivity (Fig. 4C). The integrated density of AT8 immunofluorescence in the hippocampus was decreased in some K3^RNJ1^ mice, but this reduction was not statistically significant (Fig. 4C,D). In the motor cortex, AT8 immunoreactivity showed predominantly diffuse neuropil staining, with additional intense reactivity in the soma of a subset of neurons (Fig. 4C). The integrated density of AT8 immunofluorescence in the motor cortex showed a high degree of variability within treatment arms, and group differences were not statistically significant (Fig. 4C,D). However, when inspecting the integrated AT8 fluorescence intensity against the AUC of the Rotarod Latency-to-fall for individual K3 mice in all treatment groups, we found a moderate correlation for both the hippocampus (Pearson’s correlation *r* = -0.62, *P* = 0.0018) and cortex (Pearson’s correlation *r* = -0.60, *P* = 0.0025) with Rotarod performance (AUC) (Fig. 4E). Taken together, these findings demonstrate subtle reductions in Tau pathology in whole-brain lysates of RNJ1-treated K3 mice, which were not observed in the other treatment groups.

### Quantitative proteomics and phospho-proteomics reveal reductions in abundance of human Tau and its phosphorylation in response to RNJ1 treatment

We next performed a quantitative proteomic and phosphoproteomic mass-spectrometry (MS) analysis from whole brain lysates of treated female mice as an unbiased approach to gain insight into differences between K3 and WT mice, as well as the effect of antibody treatments. To this end, we employed a TMTpro 18-plex labelling approach for quantitative analysis by tandem mass spectrometry of both tryptic peptides and phosphopeptides enriched by the “TiSH” method^26^. Given the sample-size restrictions of the labelling method, we focused the MS analysis on the WT^Veh^, K3^Veh^, K3^RNJ1^ and K3^HJ8.5^ groups, and excluded the combination group (K3^R+H^) as it had not shown synergistic effects in behavioural improvements or in Tau pathology determined by western blotting.

We found that human Tau showed a reduction of 17.7% in mean abundance in K3^RNJ1^ compared with K3^Veh^ (*P* = 0.0559), whereas K3^HJ8.5^ displayed a 9.1% reduction (*P* = 0.2124). Changes in levels of endogenous mouse Tau were more subtle, with the K3^RNJ1^ group showing a 6.7% reduction in mean abundance (*P* = 0.1960) (Fig. 4F). Interestingly, the levels of human Tau displayed a strong inverse correlation with the Rotarod performance (Pearson’s correlation *r* = -0.81, *P* = 0.0013), which was particularly pronounced for the K3^RNJ1^ group (Fig 3G). Mouse Tau levels, on the other hand, showed only a weak inverse correlation which was not statistically significant (Pearson’s correlation *r* = -0.32, *P* = 0.32). This, together with the western blot analysis, suggests that a reduction in human Tau may contribute to therapeutic efficacy in K3 mice.

We next analysed the quantitative phosphopeptide MS data for changes in the phosphorylation of individual Tau phosphosites. We edetected a total of 41 Tau phosphopeptides, comprising 27 unique phosphosites in Tau (including the well-known disease-relevant epitopes AT270, AT8, AT100, AT180, 12E8, PHF1 and PG5) (Fig 3H). The antibody treatments affected phosphorylation at these sites to various degrees. Four phosphopeptides, pS202 being part of the AT8 epitope, pS404 and pS400/pS404 being part of the PHF1 epitope, and pS409 being part of the PG5 epitope, achieved statistically significant reductions in abundance in K3^RNJ1^ compared with K3^Veh^, whereas in K3^HJ8.5^ mice, pS262 and pS356 (together forming the 12E8 epitope) were decreased. Together, our immunoblotting, histopathological and quantitative mass spectrometry analyses revealed that RNJ1-treated female K3 mice had decreased levels of total and hyperphosphorylated Tau, and that these reductions correlated with Rotarod performance.

### Quantitative proteomics reveals deregulation of proteins involved in metabolic and microtubule-related processes in K3 mice

To elucidate how RNJ1 treatment affects proteins with which Tau directly or indirectly interacts, we first aimed to delineate how the brain-wide proteome of K3 mice differs from that of wild-type mice. We found that 342 proteins out of 6,623 proteins detected in our analysis exhibited differential abundance (DA, defined as fold-change in protein levels that achieve *P* < 0.05 after false-discovery rate (FDR) correction) in K3^Veh^ relative to WT^Veh^ mice, of which 165 were decreased and 177 increased in abundance (Fig. 5A). Intriguingly, of these DA proteins, only ∼10% have been reported to interact with Tau via proximity-labelling MS^6^, encoding predominantly cytoskeletal proteins. This distribution implies that the majority of the 342 DA proteins may be affected indirectly (Fig. 5B,C).

**Figure 5.**
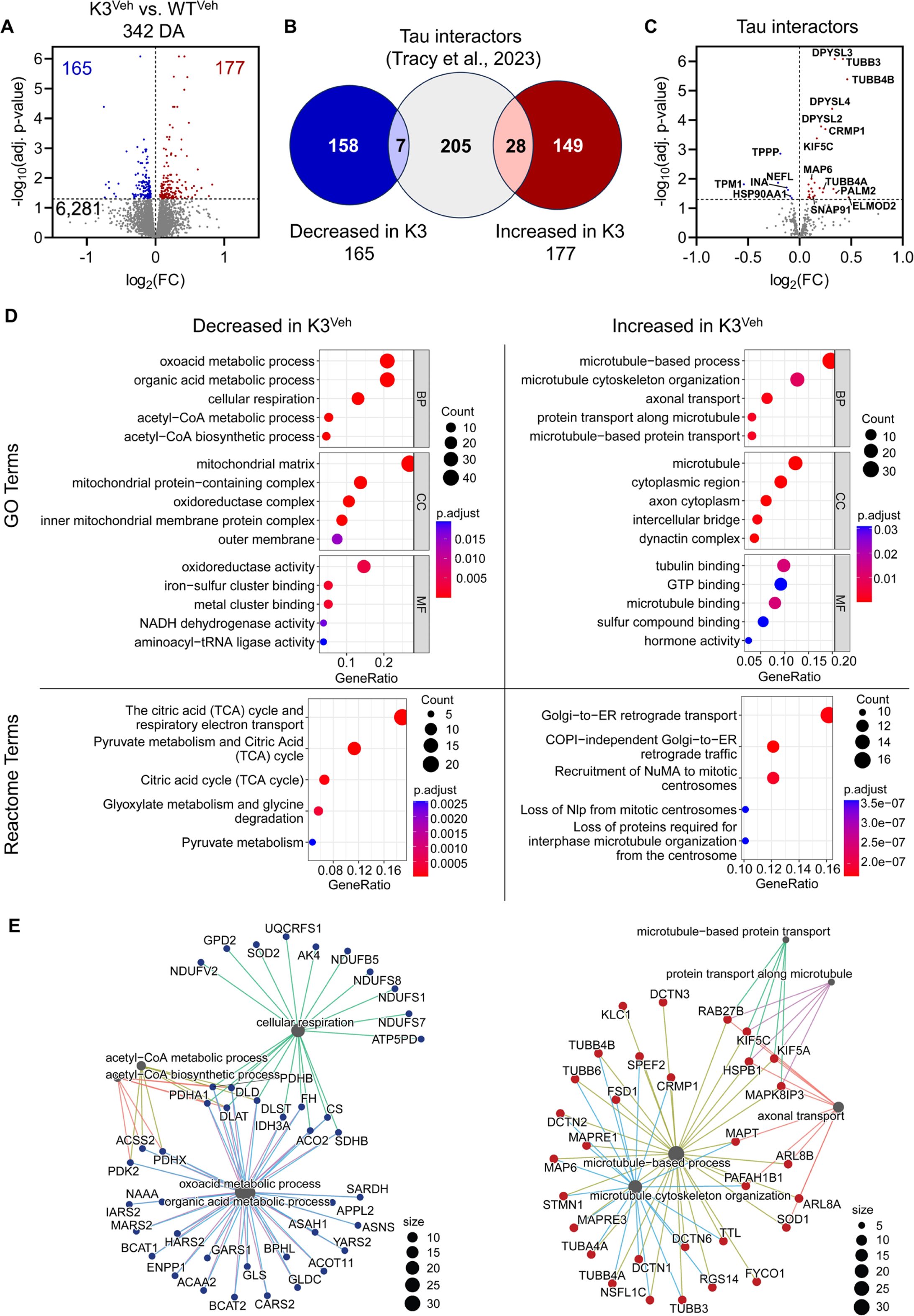
Quantitative mass spectrometry analysis shows deregulation of proteins involved in metabolic pathways and microtubule-based processes in the K3 model of tauopathy. (**A**) Volcano plot of proteins with differential abundance (DA) in K3^Veh^ relative to WT^Veh^ brains showing 342 DA proteins (165 proteins with lower and 177 with higher abundance), out of 6,623 total proteins detected in this study (6,281 with no significant changes). (**B**) Venn diagram depicting that only a small subset of the 342 proteins have recently been reported as Tau interactors via proximity-labelling MS.^6^ (**C**) Volcano plot displaying the subset of proteins in A that are reported Tau interactors. (**D**) Over-representation analysis (ORA) for proteins decreased in K3^Veh^ mice revealed Gene Ontology (GO) terms and Reactome pathways related to metabolic processes as being highly enriched, in particular proteins annotated to the terms ‘Mitochondrial Matrix’ (GO Cellular Component) and the ‘Pyruvate metabolism and citric acid (TCA) cycle’ (Reactome pathway). GO ORA for proteins with higher abundance in K3^Veh^ mice shows a strong over-representation of microtubule-based processes, with axonal transport being highly enriched. (**E**) Gene-concept networks showing DA proteins in the comparison between K3^Veh^ and WT^Veh^ associated with the most highly enriched GO terms, the latter being represented as gray nodes with connecting lines to differentially abundant proteins in K3^Veh^ mice.

We next subjected these DA proteins to a Gene Ontology (GO) and Reactome pathway over-representation analysis (ORA) to identify functional domains that are potentially affected by these DA proteins in K3 mice. By applying ORA to the protein set with lower abundance in K3 mice, several metabolic processes were found to be strongly over-represented (Fig 4D, left), with the most significantly enriched term in Reactome pathways being ‘Pyruvate metabolism and citric acid (TCA) cycle’. Conversely, in the protein set with higher abundance, over-represented terms were predominantly associated with microtubule-related processes, particularly comprising proteins with roles in axonal transport (Fig. 5D, right). Several kinesins, tubulin subtypes and dynactin subunits were upregulated, as was stathmin 1, a key regulator of microtubule dynamics, as shown by gene-concept networks of proteins annotated to enriched terms in this comparative analysis (Fig. 5E).^27,28^ Moreover, in addition to identifying small clusters of DA proteins involved in neurite outgrowth and clathrin-dependent endocytosis, STRING analysis revealed protein-protein interaction (PPI) networks of DA proteins involved in metabolic and microtubule-related processes (Supplementary Fig. 3). The pronounced decrease in proteins involved in metabolic processes, together with the increase in microtubule-related proteins, mirrors phenotypic impairments in mitochondrial function and axonal transport previously reported by us and other groups in this and other tauopathy models.^29–31^ These analyses underscore the notion that proteomic alterations in K3 mice accurately model impairments across diverse functional domains.

### Antibody treatment restores deregulated protein levels in K3 mice

We then asked whether the antibody treatment would affect the proteome (and thereby functionality) in K3 mice to restore the deregulated protein levels and associated pathways in this model. Firstly, principal component analysis (PCA) revealed that the K3^RNJ1^ mice clustered separately from the other K3 groups (K3^Veh^ and K3^HJ8.5^) (Fig. 6A). Additionally, whereas the levels of only five proteins (three up and two down) were DA in the K3^HJ8.5^ group compared to K3^Veh^, RNJ1 treatment induced DA changes to 323 proteins (186 up and 137 down) relative to the K3^Veh^ group (Fig. 6B). Together, these two analyses indicate substantial proteome-level differences in K3^RNJ1^ relative to K3^Veh^ mice.

**Figure 6.**
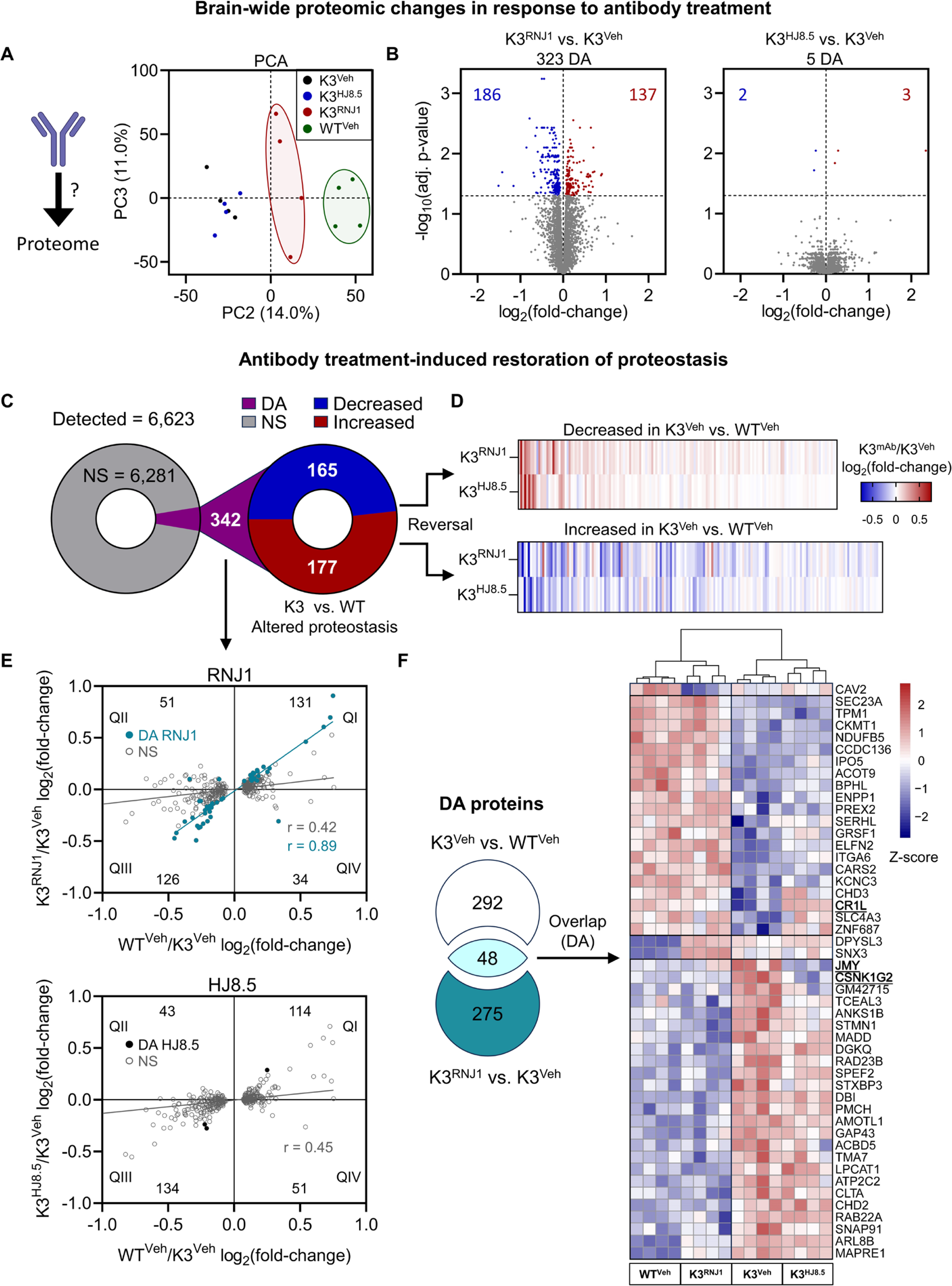
Brain-wide proteomic changes in response to antibody treatment. (**A**) Principal component analysis (PCA) reveals three clusters for PC2 and PC3, with the K3^RNJ1^ cluster shifting towards WT^Veh^. (**B**) Volcano plot for K3^RNJ1^ mice showing DA proteins (*P* < 0.05 after false-discovery rate correction), with 186 decreased and 137 increased proteins relative to K3^Veh^ (left panel). The K3^HJ8.5^ group shows higher similarity to K3^Veh^, with only 5 proteins being DA (right panel). (**C**) Analysis of therapeutic impact by focusing on the set of 342 DA proteins in K3^Veh^ compared to WT^Veh^ (pie chart and Fig. 5A). (**D**) The heatmaps represent the fold-change of proteins in the antibody-treated mice compared to K3^Veh^, with the top heatmap showing decreased and the bottom heatmap increased DA proteins in K3^Veh^ compared with WT^Veh^, showing a reversal by the antibodies towards WT^Veh^. (**E**) Scatter plot (top: RNJ1, bottom: HJ8.5) displaying the direction of change in abundance of the 342 DA proteins in WT^Veh^ and antibody-treated mice relative to K3^Veh^. The majority of the 342 proteins (∼75.1% for RNJ1; ∼72.5% for HJ8.5) fall into quadrants I and III, with quadrant I displaying proteins that are higher in WT^Veh^ and antibody-treated mice relative to K3^Veh^, and quadrant III displaying proteins that are lower in WT^Veh^ and antibody-treated mice relative to K3^Veh^, reiterating the reversal of the treatment groups towards WT. Proteins with significant fold-changes (DA) are represented as closed, and non-significant (NS) as open circles. The number of proteins falling into each quadrant is shown. (**F**) The Venn diagram represents the number of intersecting and non-intersecting proteins changed significantly (DA) in the K3^Veh^ versus WT^Veh^ and K3^RNJ^^1^ versus K3^Veh^ comparisons. The heatmap displays the row z-scores of log_2_ transformed protein abundance in the overlap, showing a striking similarity in this subset for groups WT^Veh^ and K3^RNJ1^, revealing proteins significantly affected in the K3 model, and by treatment, being restored to WT levels. Gene symbols in bold and underlined represent the corresponding DA proteins in K3^HJ8.5^ compared with K3^Veh^. Columns (individual mice) are clustered by correlation (top). Statistical comparisons of protein abundance were performed using a generalised linear model with Bayes shrinkage for group comparisons. P-values were derived from a moderated t-test and corrected for multi-hypotheses using the Benjamini and Hochberg (FDR) method.^54^

We then focused on the effects the treatment specifically had in relation to the 342 DA proteins in K3^Veh^ compared with WT^Veh^ given that this subset of proteins characterises the K3 phenotype at the proteome level (Fig. 6C). In the first instance, we analysed changes with treatment in this subset of 342 proteins without considering statistical significance, as we posited that these changes could be subtle at the individual level, but could nonetheless result in a coordinated response. Interestingly, we found that the antibody treatments increased the abundance of the majority of the proteins that were downregulated (Fig. 6D, top; Supplementary Fig. 4A), and decreased the abundance of the majority of the proteins that were upregulated in K3^Veh^ relative to WT^Veh^ (Fig. 6D, bottom; Supplementary Fig. 4B), suggesting that antibody treatment shifts the levels of deregulated proteins in K3 towards those of WT. To further understand this coordinated shift, we examined the relationship between the levels of these 342 proteins in K3^RNJ1^ and WT^Veh^ mice relative to K3^Veh^ mice (Fig. 6E, top). We first considered that antibody treatment increased or decreased protein levels with respect to K3^Veh^ mice when the fold-change values were larger or smaller than 1, respectively. We found that of the 165 proteins with decreased abundance in K3^Veh^ mice relative to WT^Veh^, 131 (∼79.4%) were shifted to higher values in K3^RNJ1^ mice (Fig 5E, top, quadrant I). Similarly, 126 (∼71.2%) of the 177 proteins with higher levels in K3^Veh^ (quadrant III) were shifted to lower abundance in K3^RNJ1^. Together, 257 of 342 (∼75.1%) DA proteins (K3^Veh^ versus WT^Veh^) were closer in abundance to WT levels after RNJ1 treatment. The same analysis indicated a similar response in K3^HJ8.5^ mice, with approximately 72.5% of these DA proteins (K3^Veh^ versus WT^Veh^) also showing changes in the same direction as WT^Veh^ (see quadrants I and III) (Fig 5E, bottom). Moreover, changes in protein levels in antibody-treated and WT^Veh^ mice relative to K3^Veh^ mice were positively correlated (Pearson’s correlation *r* = 0.42, *P* < 0.0001, for RNJ1, Fig. 6E, top; Pearson’s correlation *r* = 0.45, *P* < 0.0001, for HJ8.5, Fig. 6E, bottom).

We subsequently narrowed down the analysis to a subset of these 342 proteins that also met the criterion of being DA with antibody treatment compared with K3^Veh^. We found 48 proteins that met this criterion in the K3^RNJ1^ group (closed circles in Fig. 6E top, and Fig. 6F), and only three in the K3^HJ8.5^ group (closed circles in Fig. 6E bottom, and Fig. 6F), with the latter also featuring in the 48 proteins identified in the K3^RNJ1^ group. Remarkably, 45 of these 48 proteins in the RNJ1 group and all three in the HJ8.5 group reversed to WT levels with treatment. We again found a positive correlation between changes in protein levels in K3^RNJ1^ and WT^Veh^ mice relative to K3^Veh^ mice (Pearson’s correlation *r* = 0.89, *P* < 0.0001, closed circles in Fig. 6E top). Together, these analyses reinforce the notion that there is a restoration of protein levels in K3 towards WT following Tau antibody treatments.

In the subset of 45 overlapping DA proteins, we noticed clathrin light chain A (Clta) and clathrin coat assembly protein AP180 (Snap91), proteins involved in synaptic vesicle endocytosis, and both were decreased in abundance with RNJ1 treatment and in WT^Veh^ mice relative to K3^Veh^. To further infer the functional domains that were altered in K3 mice and potentially restored by RNJ1 treatment, we performed ORA on the overlapping DA proteins and interestingly found the highest enrichment for the GO terms ‘Presynapse’ and ‘Endocytosis’, for proteins with lower abundance in WT^Veh^ and K3^RNJ1^ relative to K3^Veh^ (Supplementary Fig. 5A,B). Furthermore, a functional PPI analysis revealed a network of 34 proteins within this subset, with six of those proteins annotated to the GO term ‘Presynapse’ and three annotated to the GO term ‘Synaptic vesicle’ (Supplementary Fig. 5C). In summary, our focused proteomic analysis indicates that both antibody treatments initiated a response to restore protein levels within the subset of proteins that characterise the K3 phenotype.

### Differential analysis of the phosphoproteome also reveals a reversion of RNJ1-treated K3 mice to wild-type levels

To understand the additional effects of antibody treatment, we performed a differential quantitative analysis of the phosphoproteome. Consistent with the proteomic analysis (Fig. 6A), a PCA of the phosphoproteome revealed three clusters: the first formed by K3^RNJ1^ mice, the second by WT^Veh^ and the third by both K3^HJ8.5^ and K3^Veh^, indicating that K3 mice have an altered phosphoproteome, which, in turn, is affected by RNJ1 treatment (Fig. 7A). We first identified 541 DA phosphopeptides in K3^Veh^ mice compared with WT^Veh^, with 283 phosphopeptides showing lower and 258 higher abundance in K3^Veh^ (Fig. 7B, left). Interestingly, 635 phosphopeptides were DA in K3^RNJ1^ compared with K3^Veh^, of which 333 displayed lower and 320 higher abundance, reflecting a shift in the phosphoproteome with RNJ1 treatment in line with the PCA analysis (Fig. 7B, middle). In contrast, no DA phosphopeptide was detected in K3^HJ8.5^ compared with K3^Veh^ (Fig. 7B, right).

**Figure 7.**
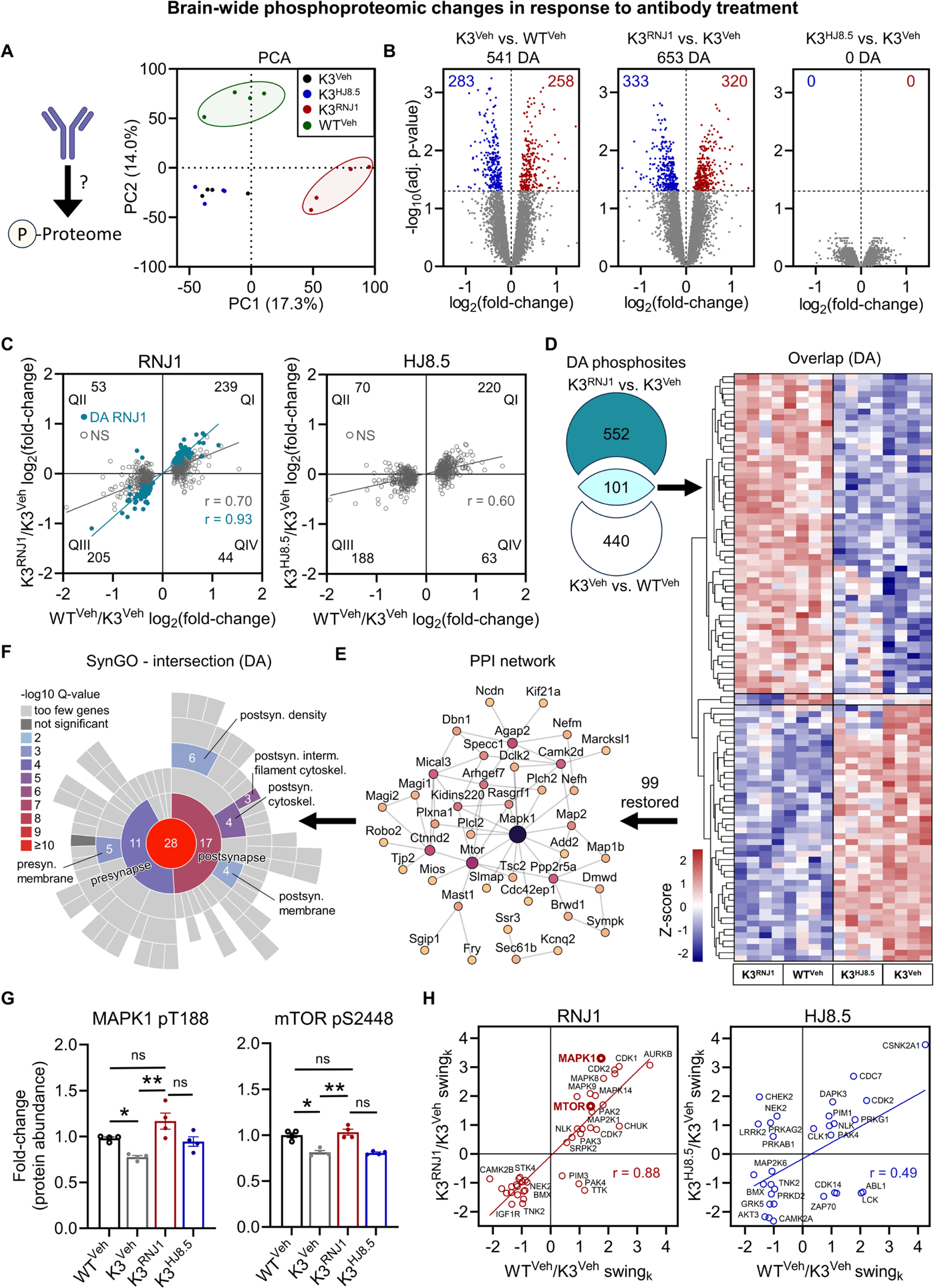
Brain-wide phosphoproteomic changes in response to antibody treatment. (**A**) Principal component analysis (PCA) of the phosphoproteome identifies three clusters, WT^Veh^, K3^RNJ1^ and one formed by K3^Veh^ and K3^HJ8.5^. (**B**) Volcano plots show differentially abundant (DA) phosphopeptides (*P* < 0.05 after false-discovery rate correction) for the K3^Veh^ versus WT^Veh^, K3^RNJ1^ versus K3^Veh^ and K3^HJ8.5^ versus K3^Veh^ comparisons. 541 phosphopeptides are DA between K3^Veh^ and WT^Veh^ mice, and 653 between K3^RNJ1^ and K3^Veh^ mice. The analysis of treatment effects on the phosphoproteome was focused on the 541 DA phosphopeptides in K3^Veh^ relative to WT^Veh^. (**C**) Scatter plots (left: RNJ1, right: HJ8.5) display the direction of change in abundance of the 541 DA phosphopeptides in WT^Veh^ and antibody-treated mice relative to K3^Veh^. The majority of the 541 DA phosphopeptides (∼82.1% for RNJ1; ∼75.4% for HJ8.5) fall into quadrants I and III, with quadrant I displaying phosphopeptides that are higher in abundance in WT^Veh^ and antibody-treated mice relative to K3^Veh^, and quadrant III those that are lower in WT^Veh^ and antibody-treated mice relative to K3^Veh^, indicating a reversal of the treatment groups towards WT. Phosphopeptides with significant fold-changes are represented as closed, and non-significant as open circles. The number of phosphopeptides falling into each quadrant and the corresponding percentage of the total of DA phosphopeptides is shown. (**D**) Venn diagram representing the number of intersecting and non-intersecting DA phosphopeptides in both K3^Veh^ versus WT^Veh^ and K3^RNJ1^ versus K3^Veh^ comparisons. The heatmap displays the row z-scores of log_2_ transformed phosphopeptide abundance in the overlap, showing striking similarity in this subset for WT^Veh^ and K3^RNJ1^, thereby revealing that phosphopeptides significantly affected in the K3 model and by treatment were restored to WT levels. (**E**) Protein-protein interaction (PPI) network of the proteins containing DA phosphopeptides shown in panel C. Node size and color relate linearly to the degree of connectivity of the node in the network. (**F**) SynGO analysis of proteins with DA phosphopeptides shown in panel C, showing enrichment of presynaptic and postsynaptic proteins. The number of proteins within the analysed set annotated to each category is shown. (**G**) S2448-phosphorylated mTOR and T188-phosphorylated MAPK1 display (different from total levels) higher abundance in K3^RNJ1^ and WT^Veh^ mice relative to K3^Veh^. Data in panel G are shown as mean ± SEM. (**H**) Network-based kinase activity prediction analysis. Scatter plots (left: RNJ1, right: HJ8.5) display the overlap of kinases with differential activity (*P* < 0.2 for swing_k_ scores) in WT^Veh^ and antibody-treated mice relative to K3^Veh^. Statistical comparisons of phosphosite abundance were performed using a generalised linear model with Bayes shrinkage for group comparisons. P-values were derived from a moderated t-test and corrected for multi-hypotheses using the Benjamini and Hochberg (FDR) method.^54^ \**P* < 0.05, \*\**P* < 0.01, \*\*\**P* < 0.001.

Akin to the proteomic analysis (Fig. 6C), we used the 541 phosphopeptides that are DA between K3^Veh^ and WT^Veh^ to determine how antibody treatments affect this subset. We first considered that antibody treatment increased or decreased phosphopeptide levels with respect to K3^Veh^ mice when the fold-change values were larger or smaller than 1, respectively (open and closed circles in Fig. 7C). We found that 205 out of 258 (∼79.4%) phosphopeptides with higher abundance in K3^Veh^ relative to WT^Veh^ were reverting towards lower abundance in the K3^RNJ1^ group (Fig. 7C, left, quadrant III). Similarly, 239 of 283 phosphopeptides (∼84.5%) with lower abundance in K3^Veh^ relative to WT^Veh^ (quadrant I) had higher abundance in K3^RNJ1^. Notably, with a total of 444 out of 541 (∼82.1%) phosphopeptides being shifted in abundance towards WT^Veh^, this degree of reversal in the phosphoproteome was even more pronounced than in the above proteomic analysis (Fig. 6D). A similar albeit less prominent reversal was observed in K3^HJ8.5^ mice, with 183 phosphopeptides being decreased and 217 increased both in K3^HJ8.5^ and WT^Veh^ relative to the K3^Veh^ group (quadrants I and III), together amounting to 408 out of 541 (∼75.4%) phosphopeptides being reverted towards WT (Fig. 7C, right).

We then examined the overlap of phosphopeptides that were statistically different in both WT^Veh^ and K3^RNJ1^ relative to K3^Veh^, and found that it comprised 101 phosphopeptides, originating from 90 proteins. Remarkably, 99 of these 101 phosphopeptides had restored levels in K3^RNJ1^ mice (Fig. 7D), mirroring the above proteomic findings. This set of 99 phosphopeptides originated from a total of 89 proteins, which we subjected to a PPI analysis which revealed a network formed by 41 proteins, with multiple functional associations between these proteins that are affected in the K3 model and are restored to WT levels upon RNJ1 treatment (Fig. 7E). Moreover, an analysis of the 89 proteins with phosphopeptides restored to WT^Veh^ levels in K3^RNJ1^ mice using the SynGO database^32^ revealed that 28 of these are proteins found in synapses, with the highest enrichment and protein count (17) found for postsynaptic proteins (Fig. 7F).

Upon examination of key proteins within the PPI network, it is noteworthy to find an increase in phosphorylation (but not total levels) of MAPK1 at threonine 188 (pT188) and of mTOR at serine 2448 (pS2448) in K3^RNJ1^ mice, with both phosphopeptides also elevated in WT^Veh^ compared with K3^Veh^ (Fig. 7G). To further explore these findings, we performed a phosphosite network-based analysis to infer the degree of activity of individual kinases, which again revealed a strong correlation between K3^RNJ1^ and WT^Veh^ mice relative to K3^Veh^ when assessing the overlap of kinases with differential activity (Pearson’s correlation *r* = 0.88, *P* < 0.0001, Fig. 7H left), whereas K3^HJ8.5^ mice showed a weaker correlation (Pearson’s correlation *r* = 0.49, *P* < 0.0001, Fig. 7H right). Furthermore, both MAPK1 and mTOR revealed increased predicted activity in K3^RNJ1^ and WT^Veh^ mice, in line with the increased phosphorylation of MAPK1 pT188 and mTOR pS2448 sites. These phosphorylation events may point towards an upregulation of the mTOR and MAPK/ERK signalling pathways, both of which are involved in a multitude of neuronal functions including synaptic plasticity.^33,34^

Together, our findings suggest that proteomic and phosphoproteomic changes may collectively reverse defects in signalling pathways and thereby contribute to improvements in functional outcomes in response to a Tau-targeting antibody therapy (Fig. 8).

**Figure 8.**
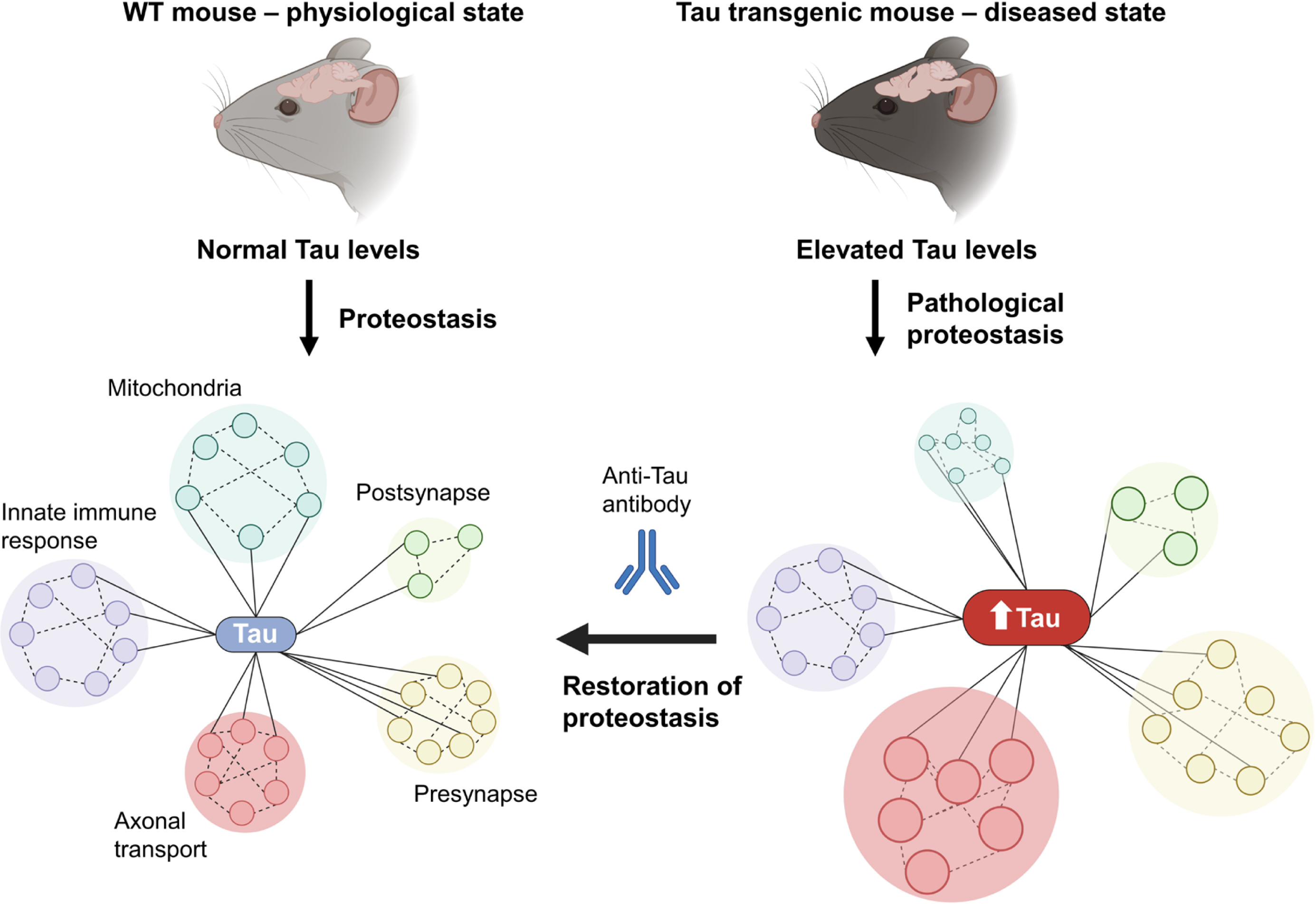
Mechanistic model for brain-wide proteomic changes induced by Tau immunotherapy. Under physiological conditions, Tau has multiple interaction partners and mediates several biological processes in neurons, including axonal transport and mitochondrial dynamics. In the K3 disease model, total Tau levels are elevated ∼3-fold relative to wild-type mice. This elevated Tau is distributed locally across different neuronal compartments, significantly impacting proteostasis as shown by our proteomic comparison between wild-type and K3, which demonstrated changes in protein abundance across functional sets. Treatment with RNJ1 reduced total Tau abundance in whole-brain extracts, which correlated with a restoration of proteostasis and improved motor function.

## Discussion

Our comparative in vitro and in vivo analysis of two Tau antibodies, one generated by us (RNJ1) and the other by Dr Holtzman and colleagues (HJ8.5, explored clinically as tilavonemab and included here for benchmarking purposes),^35–37^ revealed that these antibodies (in particular RNJ1) work towards restoring proteomic homeostasis, which we posit may serve as an additional, highly informative read-out for assessing therapeutic outcomes, rather than assessing solely Tau abundance and phosphorylation. Here, we first validated RNJ1 in the established HEK293 and a novel SH-SY5Y Tau biosensor cell system and revealed the antibody’s superior capacity in neutralising proteopathic Tau seeds derived from various tissue sources. We therefore trialed RNJ1 in the K3 mouse model of tauopathy, including three treatment arms, RNJ1 (K3^RNJ1^), HJ8.5 (K3^HJ8.5^), the two antibodies combined (K3^R+H^), and two control arms, K3 vehicle control (K3^Veh^) plus wild-type vehicle control (WT^Veh^). This demonstrated reductions in Tau pathology and improved motor functions for the RNJ1 treatment, different from the HJ8.5 and combination groups, with no synergistic effects observed for the combination treatment.

To gain a better insight into the antibodies’ impact at the brain-wide level, we conducted a comprehensive proteomic and phospho-proteomic analysis of K3 compared to WT brains, as well as the treatment arms. This revealed 342 DA proteins in K3 compared to WT mice, many of which encoded metabolic and microtubule-associated proteins, strengthening functional validations conducted in K3 mice previously and thereby underscoring the value of leveraging proteomic analyses to evaluate therapeutic effects.^23,38^ Importantly, the antibody treatments restored the levels of most proteins that were altered in K3 mice closer to those in WT mice; i.e. they induced changes towards restoring homeostasis, with RNJ1 being more effective than HJ8.5 in reverting the K3 proteomic signature towards controls.

With regard to achieving therapeutic outcomes, recent antibody developments have shifted from targeting predominantly the amino-terminus to the mid-region or phosphorylated epitopes characteristic of toxic species. Although several recent studies have demonstrated superiority of mid-region antibodies in blocking the proteopathic spread *in vitro*, this has yet to be validated in a clinical setting.^39,40^ Antibodies targeting the amino-terminus of Tau have so far suffered setbacks in clinical development, with semorinemab, gosuranemab, and tilavonemab being discontinued due to their inability to achieve their primary efficacy endpoints in phase II trials.^41–43^ Nonetheless, for lack of better antibodies, what argues for the amino-terminus as a target is this domain’s implication in pathological misfolding, thereby contributing to neurotoxicity and synaptic dysfunction.^44,45^

Here, we determined that the epitope of RNJ1 is located in the amino-terminal region and is proximal to the epitopes of HJ8.5 and RNF5, the latter being a second pan-Tau antibody generated previously in our laboratory,^20^ with the three epitopes being non-overlapping. When comparing the three antibodies’ Tau seed neutralisation capacity in vitro, RNJ1 was significantly more efficient at neutralising seeds from both the rTg4510 mouse model of tauopathy and from human AD brain tissue. While we did not explore the molecular mechanisms accounting for these differential effects, we posit that they may be related to differences in how the antibodies bind to aggregated Tau. Our results demonstrate a significantly higher reactivity of RNJ1 to aggregated Tau compared with HJ8.5 and RNF5, despite the lower affinity of RNJ1 to monomeric Tau and the proximity of their epitopes. This is in agreement with the observation that antibodies targeting amino-terminal epitopes of Tau can display differential profiles in the detection of pathological conformations of Tau.^45^ Importantly, the RNJ1 epitope (aa 9-22) identified in this study partially overlaps with the epitopes of antibodies TNT1 and TNT2, which also preferentially bind to aggregated Tau.^45^ Moreover, this epitope overlaps with the primate-specific aa 18-28 motif that mediates Tau’s interactions with several synaptic proteins,^46^ and could thereby prevent toxic synaptic Tau functions at elevated local concentrations such as those found in disease.^47^ The structural basis for the differential binding of amino-terminal antibodies to aggregated pathological Tau remains unresolved; however, our study suggests that it may have important implications for treatment efficacy and in the selection of therapeutic candidates.

Tau exists as multiple isoforms and is heavily post-translationally modified which in part affects its subcellular localisation and aggregation propensity. Given that transgenic models only partially recapitulate the human pathology, reductions of particular forms of Tau (as revealed e.g., with specific anti-phospho-Tau antibodies) may not be fully informative or predictive of functional improvements. We therefore resorted to an unbiased and more comprehensive assessment of treatment effects through quantitative proteomic and phosphoproteomic analyses. Our initial approach was to establish the set of proteins that are found with differential abundance (DA) in K3^Veh^ mice relative to WT^Veh^ mice, to then evaluate whether a response to the antibody treatments affects the levels of these proteins. Interestingly, we identified DA proteins, that are in agreement with functional read-outs we (and others) had performed previously in this and other tauopathy models, including impaired anterograde axonal transport, which affects kinesin-dependent vesicle and mitochondrial transport.^23,31^ It was noteworthy to find that a large protein subset involved in axonal transport is strongly upregulated, presumably in response to altered microtubule dynamics. Moreover, the pronounced downregulation in proteins involved in cellular respiration and metabolism mirrors reported functional mitochondrial impairments in K3 mice. We have shown previously that anterograde transport of mitochondria in K3 neurons is impaired, and we and others have shown neuronal mitochondrial dysfunction in this and additional tauopathy mouse models.^29–31^ Together, this underscores the validity of employing proteomic analyses to identify alterations in functional domains and assess the impact of therapeutic interventions.

The proteomic profiling of RNJ1-treated mice revealed notable differences to those of K3^Veh^ control and HJ8.5-treated mice. Importantly, by focusing on the set of DA proteins in the K3^Veh^ versus WT^Veh^ comparison, a key observation was most proteins had a change in DA in the direction of WT levels for both antibody treatment arms, although many of these were subtle, especially for the K3^HJ8.5^ treatment arm. Only a fraction of these were found to be statistically significant (DA) in the K3^RNJ1^ versus K3^Veh^ comparison; however, protein levels within this fraction showed a very high correlation between K3^RNJ1^ and WT^Veh^ mice, suggesting that treatment efficacy is reflected by a reversion to homeostatic protein abundance levels. It was noticeable to find an even stronger correlation in the phospho-proteomic dataset when focusing on the subset of DA phosphopeptides in K3^Veh^ relative to WT^Veh^, again for both antibody arms but with a more pronounced effect for RNJ1. The overall effects of RNJ1 on the phosphoproteome level mirrored what we had found for the proteome, i.e., the treatment eliciting a change in the same direction as WT, of which a subset was altered at a statistically significant level. Remarkably, this subset was almost completely restored to WT levels, again consistent with the changes we observed at the proteome level.

Whereas the exact mechanisms that underlie the diverse proteomic and phosphoproteomic effects were not explored in this study, our findings indicate that, by targeting Tau’s amino-terminus, both antibody treatments improved the K3 proteomic and phospho-proteomic signature. Notably, RNJ1 demonstrated a more pronounced restorative effect than HJ8.5, which likely reflects improved neuronal function and could account for the observed enhancements in motor function in our study.

Together, our data suggest that anti-Tau immunisation works towards restoring homeostasis in a tauopathy model, presenting quantitative proteomics (rather than or in addition to Tau assessments) as a viable strategy to validate therapeutic interventions in preclinical models. We foresee a more important role for omics technologies^48^ in preclinical studies and possibly, also in a clinical setting. This will be facilitated by an increased exploitation of fluid markers including plasma and cerebrospinal fluid for both diagnosis and therapeutic validation.^49–52^

## Data availability

The mass spectrometry proteomics data have been deposited to the ProteomeXchange Consortium via the PRIDE^53^ partner repository with the dataset identifier PX045799 and 10.6019/PXD045799.

Reviewer account details:

Username

Password: Dcvo7pUD

## Acknowledgements

We thank Rowan Tweedale, Dr Adam Walker and Dr. Victor Anggono for critical reading of our manuscript. We thank Keisha Roffey for help with mouse immunisations and Tishila Palliyaguru for assistance with tissue processing for histological analysis. We thank Dr. Juan Carlos Polanco for assistance with Tau seeding assays. The authors would also like to thank Dr. Martina Jones for technical assistance in performing SPR experiments and data interpretation. Image acquisition and analysis was performed at the Queensland Brain Institute’s Advance Microscopy Facility, using an automated slide scanner (Axio Imager Z2, Zeiss) with a Metafer Vslide Scanner program (Metasystems) supported by The University of Queensland through the Strategic Funding grant DVCR22052A. Animal experimentation is reported in accordance with the ARRIVE guidelines.

## Supplementary Materials for

### Supplementary Materials and Methods

#### Antibodies

Mouse monoclonal anti-Tau AT8 (pS202/pT205) (Thermo Fisher #MN1020) (WB 1:1,000, IF 1:400). Anti-Tau rabbit polyclonal antibody (Dako A0024) (WB 1:10,000). Goat anti-mouse IgG (H+L) Alexa Fluor 488 secondary antibody (Thermo Fisher #A11001) (IF 1:500). Goat-anti-mouse IR Dye^®^ 680RD (LI-COR #926-68070) (WB 1:10,000). Goat-anti-rabbit IRDye^®^ 800CW (LI-COR #926-32211) (WB 1:10,000). Polyclonal goat anti-mouse IgG (H+L) Secondary Antibody, HRP (Thermo Fisher #62-6520) (ELISA 1:5,000). The generation of the Tau antibody RNF5 was described previously ^55^.

For generation of Tau antibody HJ8.5, the VH and VL sequences as retrieved from patent WO2019180261A1 were cloned into mouse IgG2b, mouse IgG1 and kappa mAbXpress vectors, followed by large scale recombinant antibody expression and purification performed at the Queensland node of the National Biologics Facility, Australia.

#### Preparation of recombinant human Tau and single chain variable fragment (scFv) library panning

Recombinant full-length human tau 441 (hTau-441) was prepared as described previously.^56^ Purified recombinant hTau-441 was panned against the Human Single Fold scFv Libraries Tomlinson I + J. Immunotubes (Nunc) were coated with 20 µg/ml hTau-441 in PBS standing overnight at 4 °C. The next day, the tubes were washed three times with PBS and then blocked with 2% skimmed milk powder in PBS (MPBS) at room temperature for 2 h. The library (1012 phages) in skim milk powder in PBS (MPBS) was added to the coated tubes and incubated for 1 h at room temperature by rotating, followed by 1 h standing. Unbound phage was removed by washing 10 times with PBS-T (0.1% Tween 20) and then, the bound phage was eluted with 500 µl 1 mg/ml trypsin in PBS, rotating for 10 min at room temperature. E. coli XL1-blue cells (Agilent) ([OD]_600_ = 0.4) cultured in 2YT (16 g Tryptone, 10 g Yeast Extract and 5 g NaCl in 1 L distilled water) were infected with 250 µl of eluted phages and incubated in a 37 °C water bath for 30 min. To titer the phage, 10 µl of infected XL1-blue cells underwent serial dilution, and 10 µl of each dilution was then spotted on TYE plates (15 g Bacto-agar, 8 g NaCl, 10 g Tryptone, 5 g Yeast extract in 1 L distilled water) containing 100 µg/ml ampicillin and 1% glucose. The remaining infected E. coli were plated out onto a TYE plate containing 100 µg/ml ampicillin and 1% glucose. All plates were incubated overnight at 37 °C. To amplify the phage, the bacteria on the plate were collected by cell scrapping in 2 ml of 2YT, and 50 µl of the resuspended bacteria were used to inoculate 50 ml 2YT containing 100 µg/ml ampicillin plus 1% glucose. Cultures were incubated until [OD]_600_ = 0.4. KM13 helper phage (5 X 1010) was added to 10 ml of culture and incubated in a 37 °C water bath for 30 min. Cultures were then centrifuged at 3,000 x g for 10 min and pellets resuspended in 50 ml 2YT plus 100 µg/ml Ampicillin plus 50 µg/ml kanamycin plus 0.1% glucose. Cultures were incubated overnight at 30 °C shaking at 200 rpm. The phage was purified from the overnight cultures by centrifugation at 3,000 rpm for 15 min, followed by incubation with PEG/NaCl (20 % polyethylene glycol 6000, 2.5 M NaCl) for 1 h on ice. Samples were centrifuged at 3,300 x g for 30 min and pellets resuspended in 2 ml PBS ready for the next round of panning. A total of three rounds of panning was conducted. Polyclonal phage binding to hTau-441 was determined by indirect ELISA.

#### Preparation of monoclonal scFvs for ELISA

HB2151 E. coli cells ([OD]_600_ = 0.4) were inoculated with 10 µl of phage eluted from the last selection round and incubated for 30 min in a 37 °C water bath. Serial dilutions were made with 10 µl of infected HB2151 cells and 10 µl of each dilution was spotted on TYE plates containing 100 µg/ml ampicillin plus 1% glucose plates and incubated overnight at 37 °C. From the overnight plates, 20 colonies were selected for each library and inoculated in 2YT containing 100 µg/ml ampicillin plus 1% glucose in a 96 well plate and incubated overnight at 37°C at 250 rpm. Overnight cultures were then used to inoculate fresh 2YT containing 100 µg/ml ampicillin plus 0.1% glucose and incubated until 0D_600_ = 0.9. Expression of scFvs was induced with IPTG at a final concentration of 1 mM, and incubated overnight at 30 °C by shaking at 250 rpm. Plates were centrifuged at 1,800 x g for 10 min and the supernatant was collected. The supernatant was then tested for scFv binding to recombinant hTau-441 by ELISA. Positive clones were sequenced using the following primers: forward 5’-CGA CCC GCC ACC GCC GCT G-3’ and reverse 5’-CTA TGC GGC CCC ATT CA-3’.

#### Purification of scFvs and western blot analysis

RNJ1 and RNJ15 scFvs were expressed in HB2151 cells and purified from the periplasm as previously described.^57^ To further validate scFv binding to Tau, western blotting was conducted. Recombinant hTau-441 (80 ng) was electrophoresed on a 10% Tris-glycine gel (Thermo Fisher) and then transferred to a nylon membrane. The membrane was blocked in 5% skim milk in TBS containing 0.1% Tween 20 (M-TBST), and then incubated with purified scFv (1:20) in 2.5% M-TBST overnight at 4°C. ScFv binding was then detected using an anti-Myc-HRP conjugate antibody (Cell Signaling Technologies) in 2.5% MTBST for 1 hour. The blots were developed using chemiluminescence.

#### Generation of mouse IgG RNJ1

The variable domains of the clone that displayed the best binding to hTau-441, RNJ1, were separately cloned into mammalian expression vectors containing mouse IgG1 heavy and kappa light chain constant regions using InFusion cloning (National Biologics Facility, Queensland node). RNJ1 IgG was expressed in ExpiCHO cells and purified using Protein G (National Biologics Facility, Queensland node).

#### Epitope mapping of RNJ1 and confirmation of HJ8.5 epitope through indirect ELISA

Truncated human Tau forms for epitope mapping were employed as maltose binding protein (MBP) fusion proteins. For this, the corresponding truncated form was cloned into a pET His6 MBP TEV LIC plasmid (Addgene #29656) through ligation-independent cloning in frame with the amino-terminus of the maltose binding protein (MBP) cassette; then BL21(DE3) single colonies carrying the vector were used for expression and purification following the same method as for hTau-441 as described above. Nunc^®^ MicroWell™ 96 well plates were coated with the MBP-hTau fusion proteins or with hTau-441 at 10 µg/mL overnight at 4 °C and 500 RPM, followed by incubation with full-size RNJ1 and HJ8.5 at 1 µg/mL. The plates were blocked using 3% bovine serum albumin (BSA). Antibody binding was detected with an anti-mouse horseradish peroxidase (HRP) conjugate (1:10,000) followed by incubation with the HRP substrate 3,3’,5,5’-tetramethylbenzidine (Sigma-Aldrich). The reaction was stopped with 1 M HCl and absorbance measurements were taken at 450 nM using a POLARstar OPTIMA plate reader (BMG Labtech). Absorbance measurements were done in quadruplicate and blank subtracted for analysis.

For higher resolution epitope mapping of RNJ1, a library of custom synthesised peptides (Thermo Fisher) spanning the amino-terminal region of Tau containing amino acids (aa) 1-22 (peptide length 10 aa, 2 aa offset) were conjugated to BSA using EDC/NHS coupling chemistry (Thermo Fisher) and purified through size-exclusion chromatography using a Superdex 200 Increase 10/300 GL column (Cytiva) with an ÄKTApurifier chromatography system (Cytiva). The purified peptides were used to coat Nunc® MicroWell™ 96 well plates for indirect ELISA followed by incubation with RNJ1 at 1 µg/mL. Antibody detection and measurements were performed as above.

#### Determination of antibody EC50 against monomeric hTau-441

Determination of the EC_50_ of RNJ1 and HJ8.5 binding to monomeric hTau-441 was carried out through indirect ELISA as described above. ELISA plates were coated with 10 µg/mL hTau-441 and incubated with serial dilutions of RNJ1 or HJ8.5. Antibody detection and measurements were performed as above.

#### Determination of antibody binding curves against aggregated sarkosyl-insoluble Tau

For binding experiments against aggregated Tau, insoluble Tau was enriched from rTg4510 or human AD brain tissue using the previously described sarkosyl extraction method with some modifications.^58^ Briefly, for rTg4510 extracts, snap-frozen brains were homogenised in 5 volumes of RIPA buffer (Cell Signaling Technology) (20 mM Tris-HCl pH 7.5, 150 mM NaCl, 1 mM Na_2_EDTA, 1 mM EGTA, 1% NP-40, 1% sodium deoxycholate, 2.5 mM sodium pyrophosphate, 1 mM beta-glycerophosphate, 1 mM Na_3_VO_4_, 1 µg/ml leupeptin) supplemented with cOmplete^TM^ mini protease inhibitor cocktail tablets (Roche), PhosSTOP^TM^ phosphatase inhibitor tablets (Roche) and 2 mM phenylmethylsulfonyl fluoride. The homogenates were centrifuged at 20,000 x g for 20 min at 4 °C. The supernatant was transferred to a new tube and N-Lauroylsarcosine (sarkosyl) was added to a final concentration of 1% w/v. The supernatant was incubated at room temperature for 1 h with rotation, followed by centrifugation at 120,000 x g for 70 min at 22 °C using the Optima^TM^ Max-XP Ultracentrifuge (Beckman Coulter). The pellet was resuspended in ultrapure water and briefly sonicated with three 10 second pulses at 30% amplitude with a probe sonicator.

For extraction of aggregated Tau from human AD tissue, hippocampal postmortem tissue was homogenised with 10 volumes of lysis buffer containing 10 mM Tris-HCl pH 7.4, 800 mM NaCl, 5 mM EDTA, 1 mM EGTA, and 10% (w/v) sucrose supplemented with cOmplete^TM^ mini protease inhibitor cocktail tablets (Roche), PhosSTOP^TM^ phosphatase inhibitor tablets (Roche) and 2 mM phenylmethylsulfonyl fluoride. The subsequent steps were performed as for the rTg4510 tissue. In both cases, the resuspended sarkosyl insoluble pellets were purified through size-exclusion chromatography using a Superose 6 Increase 10/300 GL column (Cytiva). The void volume containing the largest fraction of seed-competent Tau was collected and concentrated with Amicon Ultra 100 kDa molecular weight cutoff filters. Protein concentrations were estimated with the BCA^TM^ Protein Assay Kit (Thermo Fisher), followed by coating of ELISA plates with sarkosyl-insoluble fractions at ∼1 µg/mL, then incubated with serial dilutions of RNJ1, HJ8.5 or RNF5. Antibody detection and measurements were performed as above.

#### Surface plasmon resonance

Surface plasmon resonance was performed on a Biacore^TM^ T200 and a Biacore^TM^ 8K system at the National Biologics Facility (NBF) at the University of Queensland. For determination of binding kinetics of RNJ1 and HJ8.5 to monomeric hTau-441, hTau-441 was immobilised onto a CM5 Series S Sensor Chip (Cytiva) using an (EDC/NHS) amine-coupling kit (Cytiva). For immobilisation, hTau-441 (40 nM in acetate buffer pH 5.5) was used with a 180 s contact time at a 5 µL/min flow rate at 25 °C. Binding affinity and binding kinetic constants were determined via multi-cycle kinetics using HPS-EP+ (Cytiva) as running buffer. Antibody injections at 3-fold increasing concentration steps (0.3-82.0 nM) (multi-cycle kinetics) were performed at a flow rate of 30 µL/min for 300 s with a subsequent dissociation time of 1,200 s. Each antibody concentration was performed in duplicate. After each injection, the sensor chip was regenerated using 10 mM glycine pH 1.7. Curve fitting and determination of kinetic constants were done using the Biacore T200 Evaluation Software.

For confirmation of epitope mapping of RNJ1 at higher resolution, the BSA-conjugated amino-terminal Tau peptides described in the ELISA section for epitope mapping, were immobilised at 40 nM onto a CM5 Series S Sensor Chip (Cytiva) using an amine-coupling kit as for hTau-441. RNJ1 injections at 3-fold increasing concentration steps (0.3-66.7 nM) (multi-cycle kinetics) were performed at a flow rate of 30 µL/min with a contact time of 180 s and a dissociation time of 600 s. Each antibody concentration was performed in duplicate. After each injection, the sensor chip was regenerated using 1 M MgCl_2_.

For assessment of antibody binding to aggregated Tau, sarkosyl-insoluble aggregated Tau extracted from brain tissue of rTg4510 mice was immobilised onto a CM5 Series S Sensor chip using amine-coupling chemistry as above. Antibody injections at 3-fold increasing concentration steps (0.3-66.7 nM) (multi-cycle kinetics) were performed at a flow rate of 30 µL/min with a contact time of 300 s and a dissociation time of 1,200 s.

#### Generation of Tau RD P301S SH-SY5Y FRET biosensor cells

Tau RD P301S SH-SY5Y FRET biosensor cells were generated using lentiviral plasmids FM5-YFP and FM5-CFP kindly provided by Prof. Marc Diamond, as described before.^59^

#### Cell culture and Tau seed neutralisation assays

Tau RD P301S HEK293 FRET biosensor cells, kindly provided by Prof. Marc Diamond, were grown in Dulbecco’s Modified Eagle Medium (DMEM) (Life Technologies) containing 10% Foetal Bovine Serum (FBS, Scientifix, SFBS-FR), 100 units/ml penicillin and 100 units/ml streptomycin (Life Technologies). SH-SY5Y FRET biosensor cells were grown in DMEM/F12 medium supplemented with 10% FBS, 15 mM HEPES, 100 units/ml penicillin and 100 units/ml streptomycin. All cells were grown in humidified incubators at 37 °C and 5% CO2.

For Tau seed neutralisation experiments with sarkosyl-insoluble Tau from rTg4510 mice, HEK293 or SH-SY5Y biosensor cells were plated at 300,000 cells/well on 24-well plates. Twenty-four hours post-plating, Tau seeds were briefly incubated with a range of antibody concentrations for 5 min in PBS, then added at 200 ng/well (sarkosyl insoluble fraction protein amount) to the supernatant. Antibody concentrations ranged from 10-300 nM when added to the wells. The cells were maintained in DMEM containing 2% FBS for 48 h, then harvested for FRET flow cytometry analysis as described previously.^60^

Seed neutralisation experiments with sarkosyl-insoluble Tau from human AD brain tissues were performed in HEK293 biosensor cells as above with the following modifications. AD sarkosyl insoluble fractions (200 ng of protein) were incubated with HJ8.5 or RNJ1 at room temperature for 5 min in 20 µL Opti-MEM. Lipofectamine^TM^ 2000 (1 µL) was added to 19 µL Opti-MEM and incubated for 5 min at room temperature. The Lipofectamine^TM^ 2000 mix was combined with the Tau-antibody mix and incubated at room temperature for 30 min. The complexes were then added dropwise to the cells (final antibody concentrations of 20 nM and 100 nM when added to the cells). The cells were maintained in DMEM containing 10% FBS for 24 h, then harvested for FRET flow cytometry analysis as above.

#### Large scale production of RNJ1 and HJ8.5 for treatment study

RNJ1 and HJ8.5 were both expressed as murine IgG1 antibodies with kappa light chains for the mouse immunisation study as we have previously obtained improved therapeutic outcomes with the IgG1 isotype.^61^ Large scale production was performed by antibody expression in ExpiCHO cells followed by Protein G affinity chromatography and SEC purification using a Superdex 200 HiLoad 26/600 column (Cytiva). The final antibody solutions were kept in a buffer containing 10 mM Tris, 100 mM NaCl, pH 7.0. This buffer was used as vehicle for groups WT^Veh^ and K3^Veh^. Antibody size and purity was confirmed by SDS-PAGE. Protein identity was confirmed by LC-MS peptide mass fingerprinting. Aggregation and purity profiles were evaluated by SE-UPLC using an Agilent Advance Bio column (300 Å, 2.7 μm, 300 x 7.8 mm). Antibody purity, size and oligomerisation profiles were further assessed by SDS capillary electrophoresis. Endotoxin levels were assessed by the Limulus amebocyte lysate (LAL) assay, with all purified samples containing <1.0 EU/mg.

#### Animals, immunisation and behavioural tests

All animal experiments were approved by the University of Queensland Animal Ethics Committee (QBI/554/17/NHMRC) and by adhering to the guidelines outlined in the Australian Code of Practice for the Care and Use of Animals for Scientific Purposes.

For the longitudinal treatment studies, 4-week-old male and female K3 mice were employed. K3 mice express human 1N4R Tau with the K369I FTD mutation under control of the mThy1.2 promoter.^62^ The mice were housed in pathogen-free cages with constant access to food and water and maintained on a 12 h light/dark cycle.^62^ Cage allocation was randomised and preserved after treatment allocation.

For treatment group allocation, forty-eight K3 mice and twelve age-matched wild-type (WT) littermates (even male-to-female ratio) were subjected to the Rotarod baseline test at 4 weeks of age, and then allocated with an equivalent average performance to five treatment groups: vehicle-treated WT mice (WT^Veh^), and K3 mice treated with vehicle (K3^Veh^), RNJ1 (K3^RNJ1^), HJ8.5 (K3^HJ8.5^), or an antibody combination (K3^R+H^). All mice handling, procedures and behavioural tests were performed in a randomised order. The treatment groups received 14 weekly intraperitoneal (IP) doses of the antibody at 50 mg/kg (25 mg/kg each for the combination) starting at 5 weeks of age. Rotarod performance was tracked longitudinally every 4 weeks, with a final test at treatment conclusion after 14 weekly treatments, amounting to a total of five Rotarod assessments.

For each Rotarod test session, the mice were habituated for a minimum of 30 min before test start. Light levels were maintained at ∼70 lx. The mice were placed on the rotating rod using a linear acceleration of 3-30 rpm over the first 90 s; the 30 rpm rotating speed was then maintained for 90 s (180 s total test time). Each mouse was tested 5 times per session and the Latency-to-fall was recorded as the longest time spent on the rod before falling out of the 5 tests. The Rotarod baseline performance was established prior to treatment start, then tracked every 4 weeks with a final test at treatment conclusion, amounting to a total of five Rotarod assessments. Rotarod tests were performed in the afternoon in a room within the Queensland Brain Institute’s animal facility in a room specifically allocated for Rotarod and grip strength assessment. Rotarod Latency-to-fall comparisons across treatment groups were performed using a two-way ANOVA with Holm-Sidak’s multiple comparisons correction. Rotarod performance throughout the treatment duration was also assessed by calculating the area under the curve (AUC) of the Latency-to-fall, and statistical comparisons were performed using a one-way ANOVA with Holm-Sidak’s multiple comparisons correction.

Grip strength of forelimbs was tested first at 4 weeks of age and once more after treatment conclusion at 18 weeks of age. For grip strength measurements, the mice were allowed to grasp a metal T-bar with their forepaws before gently pulling back from the base of the tail, ensuring that the bar and the torso remained horizontal during the pulling motion. Each mouse was tested 10 times per session and the 3 highest forces recorded were averaged as maximal grip strength.

#### Tissue processing for immunofluorescence

Forty-eight hours following completion of the behavioural tests in the treatment study, the mice were overdosed with sodium pentobarbitone and perfused transcardially with PBS. Brains were then dissected, and their hemispheres separated, removing the cerebellum. The left hemisphere was immediately snap-frozen in a dry ice bath for preservation for biochemical analysis and stored at -80°C until employed for further processing. The right hemisphere was immediately fixed in 4% PFA by immersion for 24h at 4°C, followed by dehydration by ethanol series and paraffin embedding. The paraffin-embedded hemispheres were used to obtain 7 µm sagittal sections with a rotary microtome, which were mounted onto Superfrost Plus slides (Menzel-Gläser, #SF41296SP).

#### Immunofluorescence imaging and image analysis

The paraffin embedded tissue was first dewaxed with a series of xylene and ethanol washes and rehydrated, followed by antigen retrieval in citrate buffer pH 5.8 using a microwave at 850W for 15 min. The sections were left to return to room temperature and then blocked for 1 h with blocking buffer containing 1% BSA and 20% FBS in Tris buffered saline pH 7.4 containing 0.05% Triton X-100. The sections were incubated overnight with primary antibodies diluted in blocking buffer at 4 °C. The sections were then incubated with Alexa Fluor 488-labeled goat-anti-mouse IgG secondary antibody for 4 h at room temperature and counterstained for nuclei with 0.1 µg/mL DAPI.

All images were acquired using an automated slide scanner (Axio Imager Z2, Zeiss) with a Metafer Vslide Scanner program (Metasystems) at 20x magnification. Image analysis was performed by manually drawing regions of interest corresponding to hippocampal and cortical subfields using ImageJ (FIJI). No-primary antibody control sections were used to establish mean background levels of fluorescence. The mean background values were subtracted from the experimental images prior to quantification of the integrated density within the region of interest. For all analyses, 2 to 3 sections per mice were quantified.

#### Protein extraction from whole brain lysates and western blot analysis

Frozen left hemispheres (without cerebellum) were processed with a Dounce homogeniser in six volumes of cold lysis buffer containing 50 mM HEPES pH 7.4, 2 mM EDTA, 2 mM EGTA, supplemented with cOmplete^TM^ mini protease inhibitor cocktail tablets (Roche), PhosSTOP^TM^ phosphatase inhibitor tablets (Roche) and 2 mM phenylmethylsulfonyl fluoride. At this point, a fraction of the homogenate was set aside for further processing for mass spectrometry analysis (see ‘Sample preparation for mass spectrometry’ section below). For western blot analysis, the samples were then adjusted for the lysis buffer to contain 50 mM HEPES pH 7.4, 2 mM EDTA, 2 mM EGTA, 150 mM NaCl, 1% Nonidet P-40, 0.5% sodium deoxycholate, 2.5 mM sodium pyrophosphate, 1 mM beta-glycerophosphate, 1 mM sodium orthovanadate, and 1µg/mL leupeptin. The tissue was Dounce homogenised again followed by centrifugation at 10,000g for 10 min at 4°C. The resulting supernatant was snap-frozen and stored at -80°C until further use.

Protein content in the whole brain lysates was estimated with the BCA^TM^ Protein Assay Kit (Thermo Fisher). Twenty µg of protein was then boiled for 5 min in Laemmli sample buffer and resolved by SDS-PAGE on a 4-15% Criterion^TM^ TGX^TM^ precast gel (Bio-Rad). Electrophoresed proteins were transferred to Immun-Blot Low Fluorescence PVDF membranes (Bio-Rad) using the Trans-Blot Turbo Transfer System (Bio-Rad), followed by staining with REVERT^TM^ 700 Total Protein Stain (LI-COR) as per the manufacturer’s instructions. The membranes were then washed and blocked for 1 h at room temperature with blocking buffer (5% BSA in TBS), followed by overnight incubation at room temperature with primary antibody in blocking buffer. The membranes were washed and probed with fluorescent secondary antibodies for 1 h at room temperature. After washing, the membranes were imaged in the 680 nm and 800 nm channels with the Odyssey FC Imaging System (LI-COR). Fluorescent signal quantification and analysis was performed with the Image Studio^TM^ software (LI-COR). Background subtracted values were normalised to total protein (REVERT^TM^ 700) signal.

#### Sample preparation for mass spectrometry

To the homogenate prepared for mass spectrometry analysis (see ‘Protein extraction from whole brain lysates and western blot analysis’ section above), an aliquot of a 10% sodium dodecyl sulfate (SDS) stock solution was added to obtain a final concentration of 2% SDS, which was heated at 85 °C for 10 min. The sample was then stored at -80 °C until use.

The samples were thawed and reduced at 85 °C for 10 min by adding tris(2-carboxyethyl)phosphine to a final concentration of 10 mM. 2-iodoacetamide was then added to a working concentration of 20 mM and the sample incubated at 23 °C for 30 min, followed by precipitating the proteins in the samples using the chloroform-methanol method.^63^

The precipitate was dried and dissolved in 7.8 M Urea/100 mM HEPES pH 8.0/LysC solution, adding a total of 5 µg LysC to each sample. LysC digestion was performed for 12 h at 28 °C. This was followed by two trypsin digestions at 28 °C for 8 h each, using 5 µg of trypsin (TrypZean, Sigma) for each reaction. 250 µg of the tryptic peptides from each sample were made up to an equal concentration across all samples in 100 mM HEPES pH 8.0 and then labelled with TMTpro 18-plex reagents (Thermo Fisher) according to the manufacturer’s instructions. The reaction was quenched with 5% hydroxylamine for 10 min, after confirming that efficient labelling occurred by a brief LC-MS/MS analysis of a small fraction of each sample. The samples were pooled and dried to a small volume. The combined sample was desalted using solid phase extraction cartridges (Sep-Pak Vac 3cc t18, Waters). Phosphopeptides were then enriched following the “TiSH” method.^64^

The phosphopeptide-enriched fraction and approximately 200 µg of phosphopeptide de-enriched sample were fractionated by hydrophilic interaction chromatography (HILIC). A Dionex Ultimate 3000 HPLC system was used with a 250 mm long and 1 mm inside diameter TSKgel Amide-80 column (Tosoh Biosciences). The HILIC gradient that was used consisted of 90% acetonitrile, 0.1% TFA (buffer A) and a solution of 0.1% TFA (buffer B). The sample was injected into a 250 µL sample loop. The flow rate was 60 µL/min in Buffer A for 10 min to load the sample. The gradient was from 100% to 60% Buffer A for 35 min at a flow rate of 50 µL/min. The fractions were collected into a 96-well plate using a Probot (LC packings) at 30-second intervals, monitored by UV absorbance at 214 nm. The UV signal was used as a guide to combine selected fractions into similar amounts of peptide.

#### LC-MS/MS analysis

LC-MS/MS was performed using a Dionex UltiMate 3000 RSLC nano system and a Q Exactive Plus hybrid quadrupole-orbitrap mass spectrometer (Thermo Fisher Scientific). Each HILIC phosphopeptide-enriched fraction was loaded directly onto an in-house 300 mm long column with a 0.075 mm inside diameter packed with ReproSil Pur C18 AQ 1.9 µm resin (Dr Maisch, Germany). The column was heated to 50 °C using a column oven (PRSO-V1, Sonation Lab Solutions) integrated with the nano flex ion source with an electrospray operating at 2.3 kV. The S lens radio frequency level was 50 and the capillary temperature was 250 °C.

Twenty-six phosphopeptide fractions were analysed using a 120 min LC-MS/MS method. An aliquot of 5 µL of each fraction was injected into a 20 µL loop and loaded onto the column in 99% reverse phase buffer A (solution of 0.1% formic acid) and 1% buffer B (solution of 0.1% formic acid, 90% acetonitrile) for 25 min at 300 nL/min. The gradient, at 250 nL/min, was from 1% buffer B to 5% buffer B in 1 min, to 25% buffer B in 74 min, to 35% buffer B in 8 min, to 99% buffer B in 1 min, held at 99% buffer B for 2 min, to 99% buffer A in 1 min and held for 8 min as the flow rate ramped up to 300 nL/min. MS acquisition was performed for the entire 120 min. All samples and fractions were analysed using data-dependent acquisition LC-MS/MS. The MS scans were at a resolution of 70,000 with an automatic gain control target of 1,000,000 for a maximum ion time of 100 ms from 375 to 1500 m/z. The MS/MS scans were at a resolution of 35,000 with an automatic gain control target of 200,000 and maximum ion time of 115 ms. The loop count was 12, the isolation window was 1.1 m/z, the first mass was fixed at 120 m/z and the normalised collision energy was 34. Singly charged ions and those with charge greater than 8 were excluded from MS/MS and dynamic exclusion was for 35 s.

Twenty-seven non-phosphopeptide fractions were analysed using a 140 min LC-MS/MS method. An aliquot of 3.5 µL of each fraction was injected into a 20 µL loop and loaded onto the column in 99% reverse phase buffer A and 1% buffer B for 17.5 min at 300 nL/min. The gradient, at 250 nL/min, was from 1% buffer B to 6% buffer B in 1 min, to 28% buffer B in 101.5 min, to 35% buffer B in 8 min, to 99% buffer B in 1 min, held at 99% buffer for 2 min, to 99% buffer A in 1 min and held for 8 min as the flow rate ramped up to 300 nL/min. MS acquisition was performed for the entire 140 min. All samples and fractions were analysed using data-dependent acquisition LC-MS/MS. The MS scans were at a resolution of 70,000 with an automatic gain control target of 1,000,000 for a maximum ion time of 100 ms from 375 to 1,500 m/z. The MS/MS scans were at a resolution of 35,000 with an automatic gain control target of 200,000 and maximum ion time of 100 ms. The loop count was 12, the isolation window was 1.1 ms, the first mass was fixed at 120 m/z and the normalised collision energy was 31. Singly charged ions and those with charge greater than 8 were excluded from MS/MS and dynamic exclusion was for 37 s.

#### MS data processing

The raw LC-MS/MS data was processed with MaxQuant v1.6.7.0. Variable modifications were oxidation (M), acetyl (protein N-terminus), deamidation (N and Q) and phosphorylation (S, T, and Y). Carbamidomethyl (C) was a fixed modification. Digestion was set to trypsin/P with a maximum of 3 missed cleavages. Minimum reporter peptide ion fraction was 0.6. The *Mus musculus* UniProt reference proteome was used with canonical and isoform sequences (downloaded Aug 11 2022 with 55,315 entries and 21,984 genes). A fasta file containing the human Tau K369I transgenic protein (P10636-7 Uniprot entry for human Tau 1N4R sequence with the K369I mutation) was also used, as well as the inbuilt MaxQuant contaminants fasta file. The minimum peptide length was 6 and maximum peptide mass was 6000, the peptide and protein false discovery rates were set at 1%. The minimum score for modified peptides was 40. All modified peptides and counterpart non-modified peptides were excluded from protein quantification. All other settings were default.

### MaxQuant data processing of the proteome and phosphoproteome of the antibody treatment study

After processing spectra with MaxQuant, output “evidence.txt” and “proteinGroups.txt” files were further processed for proteome and phosphoproteome analysis, respectively (all analyses were performed in R version 4.05).^65^

#### Proteome

After protein IDs were annotated to gene symbols from the corresponding protein fasta file used for the MaxQuant search, intensity values of complete data (i.e. no missing values for any replicates) was log_2_ transformed and quantile normalised,^66^ followed by surrogate variable analysis for removal of unwanted sources of variation ^67^ using a predefined model describing five groups: (1) WT^Veh^, (2) K3^Veh^, (3) K3^RNJ1^, (4) K3^HJ8.5^, and (5) K3^R+H^. Hierarchical clustering and PCA were used to guide surrogate variable analysis correction. For analysis of Tau abundance, due to homology between the human Tau K369I transgene and the endogenous mouse Tau, peptide reassignment was performed using only the unique peptides for each isoform. For differential protein analysis, a generalised linear model with Bayes shrinkage, as implemented in limma version 3.46.0,^68^ was fit and each of the sample groups contrasted. P-values were derived from a moderated t-test and corrected for multi-hypotheses using the Benjamini and Hochberg (FDR) method.^69^ DA proteins were defined as those with p < 0.05 after FDR correction.

#### Phosphoproteome

Further processing of MaxQuant evidence files of phosphopeptides was performed as described previously.^64,70,71^ Briefly, the same fasta file used for MaxQuant data processing was employed to map phosphopeptides identified in the MaxQuant search. Peptide sequences with a window of +/- 7AA around each phosphorylation site were subsequently annotated as previously described. Only phosphopeptides with a site localisation score ≥ 0.75 (class I) for a minimum of one site within a phosphopeptide detected across all the samples were retained for further analysis. In the case of multi-phosphorylated peptides, additional phosphosites within the same peptides containing a class I site required a score > 0.5. For some phosphopeptides, multiple measurements were obtained – in these cases the median log_2_ transformed intensity of all detections was recorded. Phosphopeptides mapped to more than one unique protein were assigned to unique multi-mapped protein identifiers. Only measurements with no missing values for any of the replicates were considered. Intensity values were quantile normalised followed by surrogate variable analysis for removal of unwanted sources of variation^67^ using a predefined model describing five groups: (1) WT^Veh^, (2) K3^Veh^, (3) K3^RNJ1^, (4) K3^HJ8.5^, and (5) K3^R+H^. Hierarchical clustering and principal component analysis were used to guide surrogate variable analysis correction. We employed a generalised linear model with Bayes shrinkage (limma version 3.46.0)^68^ for differential activity. Reported p-values (from a moderated t-test) were corrected for multiple hypotheses testing with the Benjamini and Hochberg (FDR) method.^69^ DA phosphopeptides were defined as those with p < 0.05 after FDR correction.

#### Protein kinase activity prediction

Protein kinase activity prediction was performed using the network theoretic approach, KinSwing, as described previously,^72^ with the additional implementation of significance scoring of kinase activity. First, kinase matches are modelled as position weight matrices of known mammalian kinase:substrate sequences and their probabilistic matches to each phosphorylated peptide identified. Kinase activity or *swing_k_*^72^ is then determined by the weighted local network connectivity of kinase:substrate interactions, as the absolute proportion of significant positive and negative substrates that have significant fold changes (p ≤ 0.05), weighted for edge count and substrate count of sequences used to build the kinase position weight matrix model and z-transformed. Significance of inferred kinase activity *swing_k_* is then determined by permutating network kinase node labels, *K*, to substrates, *S*, of the total network, *M_ks_*. Thus the probability of observing *swing_k_* is conditional on this permuted reference distribution, of size, *N* (1,000 permutations), and computed for each tail of the distribution, positive and negative *swing_k_* scores, for reporting of kinase activity p-values.

#### Other bioinformatic and statistical analysis

The sample size for the treatment study was established based on previous studies to achieve adequate power for behavioural improvements. Gene set over-representation analysis with GO, Reactome and KEGG pathway knowledgebases was performed with R package *clusterProfiler* version 4.6.2^73^ or with Metascape,^74^ using DA proteins with increased or decreased differential abundance as foreground and the complete list of proteins detected in this study as a custom background. Heatmaps were generated with the R package *pheatmap* version 1.0.12 (https://CRAN.R-project.org/package=pheatmap) or with Prism 9 (GraphPad Software). For proteins with differential abundance in K3^Veh^ mice compared with WT^Veh^, a functional protein-protein interaction network with a STRING confidence cutoff = 0.7 was represented in Cytoscape (v3.9.1). For proteins with restored phosphosite abundance in K3^RNJ1^ and WT^Veh^ compared with K3^Veh^, a “Physical (Core)” protein-protein interaction network analysis was performed with Metascape and represented in Cytoscape. Analysis of synaptic annotations and term enrichment for proteins with DA phosphopeptides was performed with the SynGO database. For the analysis of antibody treatment effects on the phosphoproteome, we excluded phosphopeptides in cases where the phosphopeptide and the protein it originates from were both differentially abundant in the same direction within the corresponding comparison. This was done to avoid capturing effects on the phosphoproteome that predominantly reflect what had already been observed regarding differential protein abundance in the proteome analysis.

All other statistical analyses were performed in Prism 9 (GraphPad Software). Data are represented as mean ± SEM throughout the study unless otherwise stated. For multiple comparisons, one-way ANOVA with Holm-Sidak’s correction was performed unless otherwise stated. For comparison of two groups, unpaired t-tests were used unless stated differently.

## Supplementary Figure Legends

**Supplementary Fig. 1.**
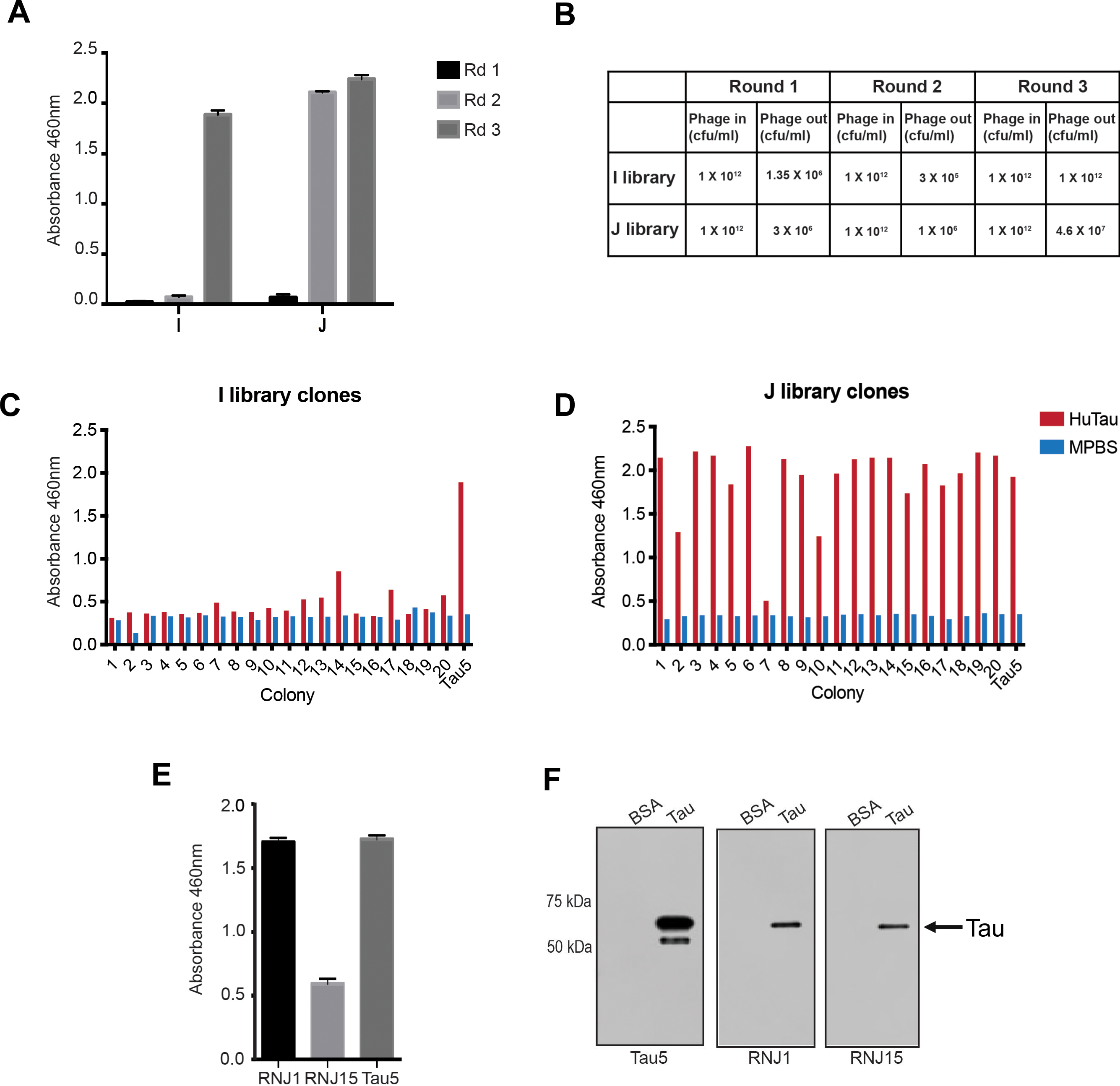
Isolation of human Tau binders after panning of Tomlinson I + J libraries. (**A**) Three successive rounds of panning were conducted against full-length human Tau (hTau-441). ELISA of the polyclonal phage from each round of panning, and (**B**) Phage colony forming units (cfu) counts from each round of panning, revealing an enrichment in positive binders for hTau-441 from each library. ELISA of isolated scFvs following the third round of panning from the I library (**C**) and the J library (**D**) tested against hTau-441 and control, skim milk powder in PBS (MPBS). (**E**) ELISA of isolated scFv clones RNJ1 and RNJ15 against recombinant hTau-441 and compared to the Tau monoclonal antibody, Tau5. (**F**) scFv clones RNJ1 and RNJ15 demonstrate specific binding to hTau-441 but not control protein, BSA, as shown by western blotting.

**Supplementary Fig. 2.**
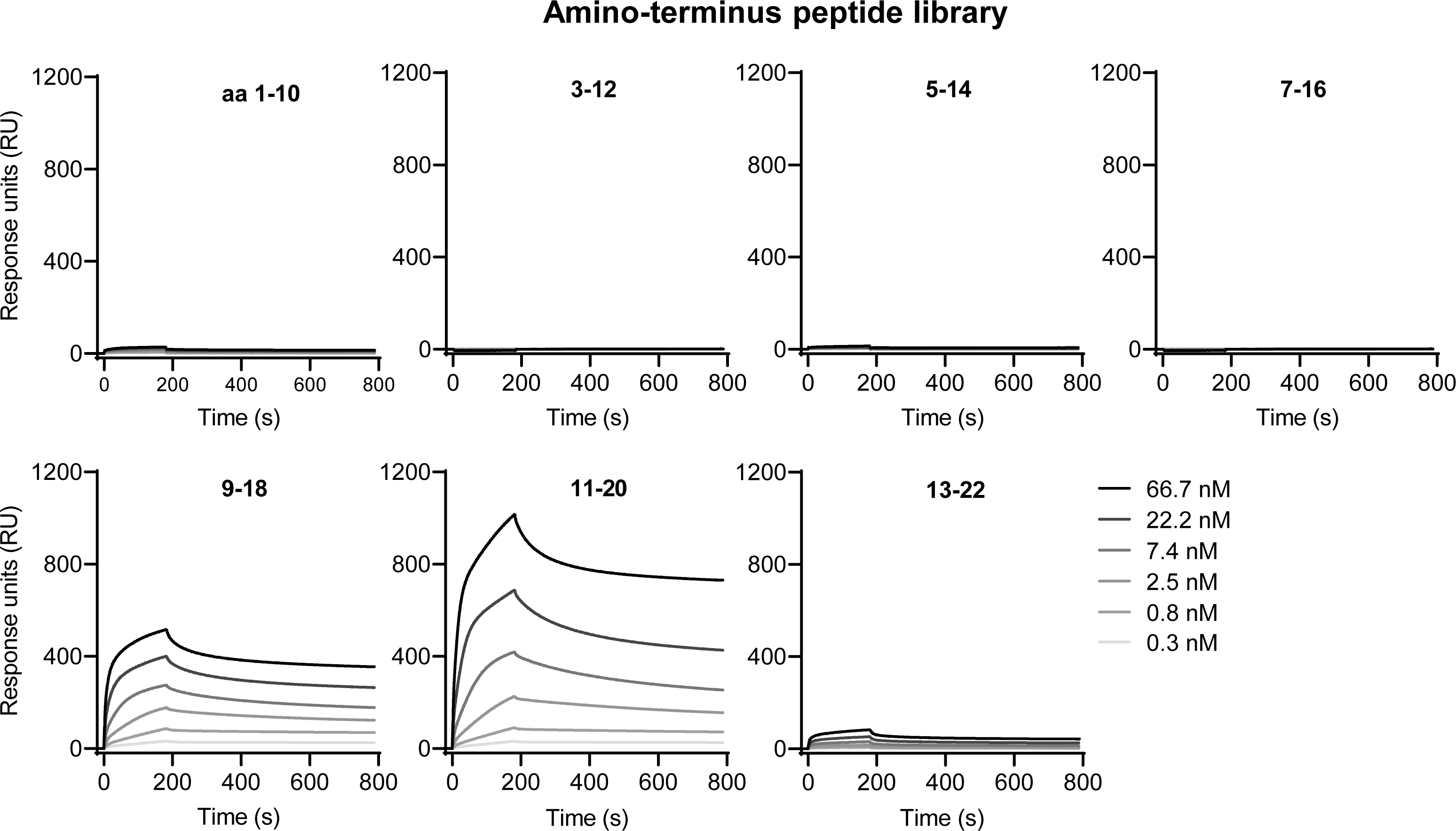
Binding of RNJ1 to the amino-terminus of Tau determined by surface plasmon resonance. Peptides from a synthetic library spanning amino-acids (aa) 1-22 of human Tau were first conjugated to BSA and then used as ligands immobilised on SPR sensor chips. RNJ1 was then used as analyte to detect binding to the different peptides.

**Supplementary Fig. 3.**
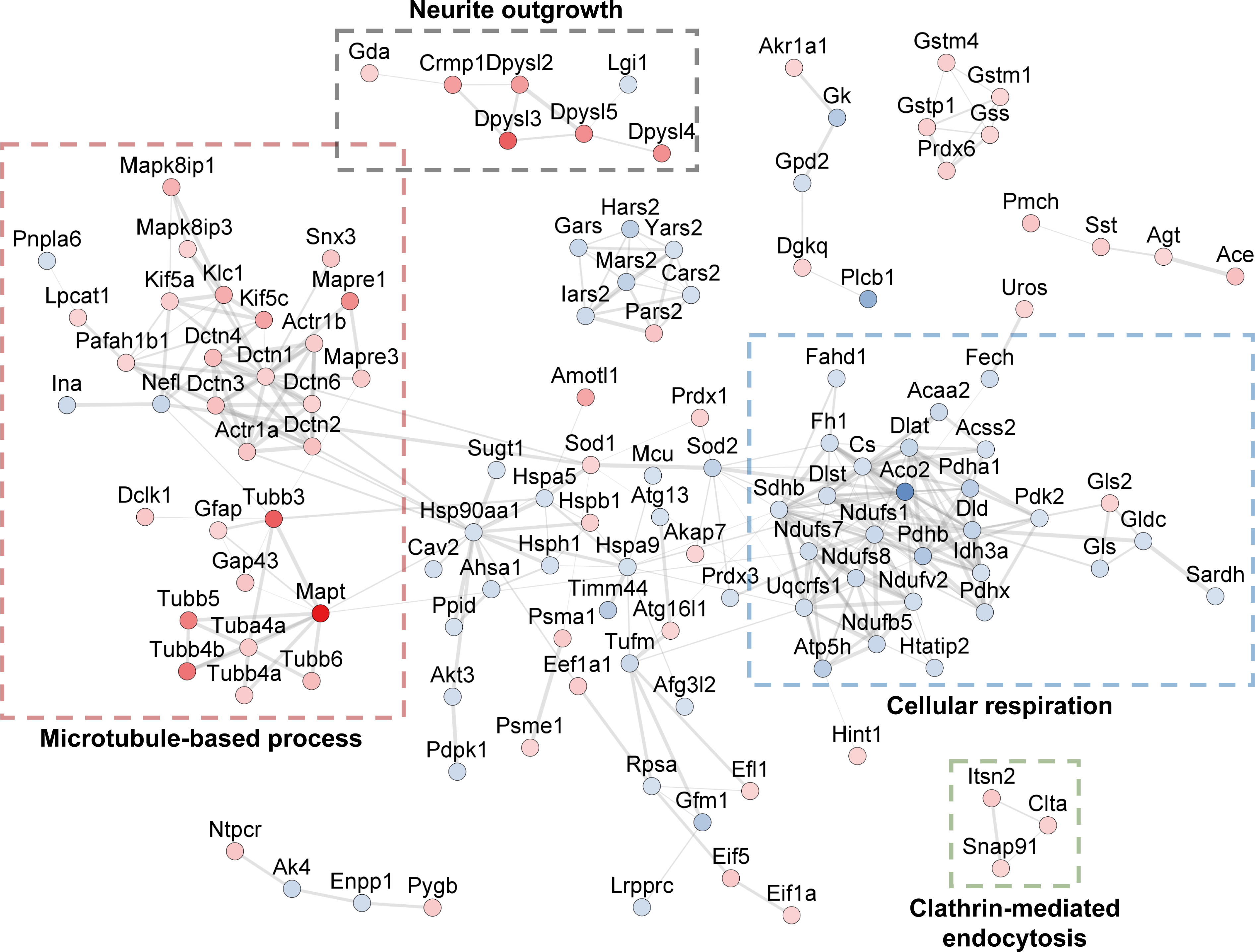
Functional protein-protein interaction network of differentially abundant proteins in K3^Veh^ relative to WT^Veh^ mice. Functional STRING network using a confidence cutoff = 0.7 for all differentially abundant (DA) proteins in K3^Veh^ relative to WT^Veh^ mice. Clusters formed by >3 proteins are displayed. Protein circles are color-coded with a linear continuous mapping of their t-statistic in the comparison between K3^Veh^ and WT^Veh^ mice. Line widths reflect the confidence of the association. Dashed rectangles highlight clusters where the majority of proteins are associated with the biological process listed.

**Supplementary Fig. 4.**
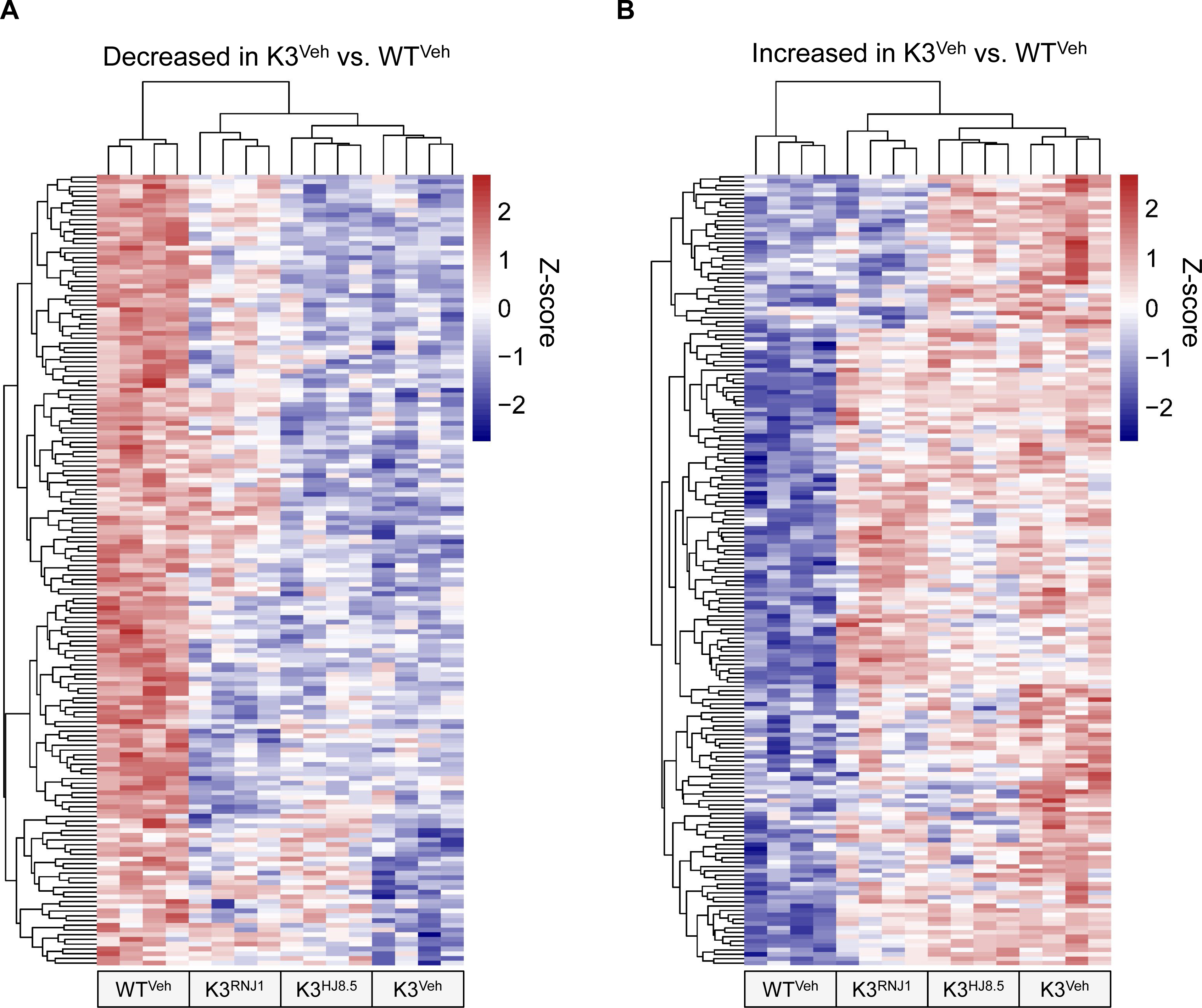
Heatmaps of all differentially abundant proteins in K3^Veh^ relative to the WT^Veh^ group. (**A**) Heatmap of all proteins increased in K3^Veh^ relative to WT^Veh^ showing row z-score values for treatment groups WT^Veh^, K3^RNJ1^, K3^HJ8.5^ and K3^Veh^. (**B**) Corresponding heatmap of all proteins decreased in K3^Veh^ relative to WT^Veh^. Both heatmaps shown as row z-scores of the log_2_ transformed protein abundance values. Columns and rows were clustered by correlation.

**Supplementary Fig. 5.**
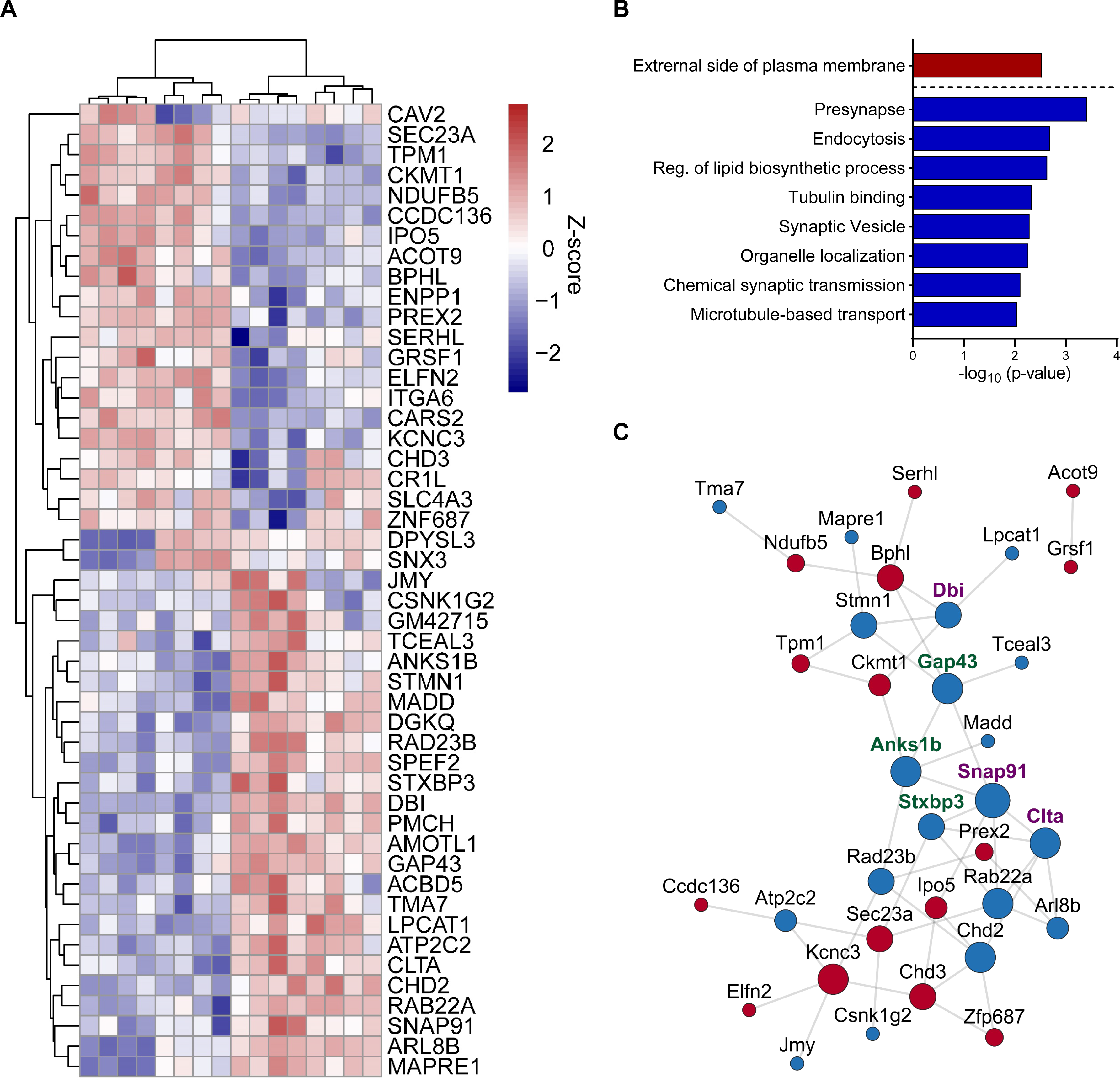
Analysis of the effects of antibody treatment on proteins that are altered in K3^Veh^ mice. (**A**) Heatmap of proteins that are differentially abundance (DA) in WT^Veh^ and K3^RNJ1^ compared with K3^Veh^ (see Fig. 6F), showing restoration of wild-type protein levels for K3^RNJ1^ for 45 of 48 proteins. The heatmap displays the row z-scores of the log_2_ transformed protein abundance values. (**B**) Over-representation analysis of DA proteins in the overlap shown in panel A, with enriched terms for proteins with increased abundance in WT^Veh^ and K3^RNJ1^ mice shown in red, and for proteins decreased in abundance in WT^Veh^ and K3^RNJ1^ shown in blue. (**C**) Functional PPI of proteins from panel A. The node size is linearly encoded by connectivity. The node color describes the direction of protein abundance fold-change in WT^Veh^ and K3^RNJ1^ relative to K3^Veh^. Green labels display proteins annotated to the GO term ‘Presynapse’. Purple labels display proteins annotated to both GO terms ‘Presynapse’ and ‘Synaptic vesicle’.

**Supplementary data file 1**. Raw data for all experiments where n < 20

## References

1. Polanco JC, Li C, Bodea LG, Martinez-Marmol R, Meunier FA, Götz J. Amyloid-β and tau complexity - towards improved biomarkers and targeted therapies. Nat Rev Neurol. 2018;14(1):22–39.

2. Frost B, Hemberg M, Lewis J, Feany MB. Tau promotes neurodegeneration through global chromatin relaxation. Nat Neurosci. 2014;17(3):357–66.

3. Udeochu JC, Amin S, Huang Y, et al. Tau activation of microglial cGAS-IFN reduces MEF2C-mediated cognitive resilience. Nat Neurosci. 2023;26(5):737–750.

4. Conze C, Rierola M, Trushina NI, et al. Caspase-cleaved tau is senescence-associated and induces a toxic gain of function by putting a brake on axonal transport. Mol Psychiatry. 2022;27(7):3010–3023.

5. Eteläinen TS, Silva MC, Uhari-Väänänen JK, et al. A prolyl oligopeptidase inhibitor reduces tau pathology in cellular models and in mice with tauopathy. Sci Transl Med. 2023;15(691):eabq2915.

6. Tracy TE, Madero-Pérez J, Swaney DL, et al. Tau interactome maps synaptic and mitochondrial processes associated with neurodegeneration. Cell. 2022;185(4):712–728.e14.

7. Cummings J, Zhou Y, Lee G, Zhong K, Fonseca J, Cheng F. Alzheimer’s disease drug development pipeline: 2023. Alzheimers Dement (N Y). 2023;9(2):e12385.

8. Long JM, Holtzman DM. Alzheimer Disease: An Update on Pathobiology and Treatment Strategies. Cell. 2019;179(2):312–339.

9. Perneczky R, Jessen F, Grimmer T, et al. Anti-amyloid antibody therapies in Alzheimer’s disease. Brain. 2023;146(3):842–849.

10. Larkin HD. Lecanemab Gains FDA Approval for Early Alzheimer Disease. JAMA. 2023;329(5):363.

11. Sims JR, Zimmer JA, Evans CD, et al. Donanemab in Early Symptomatic Alzheimer Disease: The TRAILBLAZER-ALZ 2 Randomized Clinical Trial. JAMA. 2023;330(6):512–527.

12. Dujardin S, Colin M, Buée L. Invited review: Animal models of tauopathies and their implications for research/translation into the clinic. Neuropathol Appl Neurobiol. 2015;41(1):59–80.

13. Yanamandra K, Jiang H, Mahan TE, et al. Anti-tau antibody reduces insoluble tau and decreases brain atrophy. Ann Clin Transl Neurol. 2015;2(3):278–88.

14. Furman JL, Diamond MI. FRET and Flow Cytometry Assays to Measure Proteopathic Seeding Activity in Biological Samples. Methods Mol Biol. 2017;1523:349–359.

15. Johnson ECB, Carter EK, Dammer EB, et al. Large-scale deep multi-layer analysis of Alzheimer’s disease brain reveals strong proteomic disease-related changes not observed at the RNA level. Nat Neurosci. 2022;25(2):213–225.

16. Askenazi M, Kavanagh T, Pires G, Ueberheide B, Wisniewski T, Drummond E. Compilation of reported protein changes in the brain in Alzheimer’s disease. Nat Commun. 2023;14(1):4466.

17. Bhaskar K, Hobbs GA, Yen SH, Lee G. Tyrosine phosphorylation of tau accompanies disease progression in transgenic mouse models of tauopathy. Neuropathol Appl Neurobiol. 2010;36(6):462–77.

18. Li C, Götz J. Somatodendritic accumulation of Tau in Alzheimer’s disease is promoted by Fyn-mediated local protein translation. EMBO J. 2017;36(21):3120–3138.

19. Santacruz K, Lewis J, Spires T, et al. Tau suppression in a neurodegenerative mouse model improves memory function. Science. 2005;309(5733):476–81.

20. Bajracharya R, Cruz E, Götz J, Nisbet RM. Ultrasound-mediated delivery of novel tau-specific monoclonal antibody enhances brain uptake but not therapeutic efficacy. J Control Release. 2022;349:634–648.

21. Dujardin S, Commins C, Lathuiliere A, et al. Tau molecular diversity contributes to clinical heterogeneity in Alzheimer’s disease. Nat Med. 2020;26(8):1256–1263.

22. Holmes BB, Furman JL, Mahan TE, et al. Proteopathic tau seeding predicts tauopathy in vivo. Proc Natl Acad Sci USA. 2014;111(41):E4376–85.

23. Ittner LM, Fath T, Ke YD, et al. Parkinsonism and impaired axonal transport in a mouse model of frontotemporal dementia. Proc Natl Acad Sci USA. 2008;105(41):15997–6002.

24. Pandit R, Leinenga G, Götz J. Repeated ultrasound treatment of tau transgenic mice clears neuronal tau by autophagy and improves behavioral functions. Theranostics. 2019;9(13):3754–3767.

25. Noble W, Hanger DP, Miller CC, Lovestone S. The importance of tau phosphorylation for neurodegenerative diseases. Front Neurol. 2013;4:83.

26. Engholm-Keller K, Larsen MR. Improving the Phosphoproteome Coverage for Limited Sample Amounts Using TiO2-SIMAC-HILIC (TiSH) Phosphopeptide Enrichment and Fractionation. Methods Mol Biol. 2016;1355:161–77.

27. Lyu J, Yang EJ, Zhang B, et al. Synthetic lethality of RB1 and aurora A is driven by stathmin-mediated disruption of microtubule dynamics. Nat Commun. 2020;11(1):5105.

28. Yadav P, Selvaraj BT, Bender FL, et al. Neurofilament depletion improves microtubule dynamics via modulation of Stat3/stathmin signaling. Acta Neuropathol. 2016;132(1):93–110.

29. David DC, Hauptmann S, Scherping I, et al. Proteomic and functional analyses reveal a mitochondrial dysfunction in P301L tau transgenic mice. J Biol Chem. 2005;280(25):23802–14.

30. Cummins N, Tweedie A, Zuryn S, Bertran-Gonzalez J, Götz J. Disease-associated tau impairs mitophagy by inhibiting Parkin translocation to mitochondria. EMBO J. 2019;38(3):e99360.

31. DuBoff B, Götz J, Feany MB. Tau promotes neurodegeneration via DRP1 mislocalization in vivo. Neuron. 2012;75(4):618–32.

32. Koopmans F, van Nierop P, Andres-Alonso M, et al. SynGO: An Evidence-Based, Expert-Curated Knowledge Base for the Synapse. Neuron. 2019;103(2):217–234.e4.

33. Eng AG, Kelver DA, Hedrick TP, Swanson GT. Transduction of group I mGluR-mediated synaptic plasticity by β-arrestin2 signalling. Nat Commun. 2016;7:13571.

34. Tang SJ, Reis G, Kang H, Gingras AC, Sonenberg N, Schuman EM. A rapamycin-sensitive signaling pathway contributes to long-term synaptic plasticity in the hippocampus. Proc Natl Acad Sci USA. 2002;99(1):467–72.

35. Yanamandra K, Kfoury N, Jiang H, et al. Anti-tau antibodies that block tau aggregate seeding in vitro markedly decrease pathology and improve cognition in vivo. Neuron. 2013;80(2):402–414.

36. Ising C, Gallardo G, Leyns CEG, et al. AAV-mediated expression of anti-tau scFvs decreases tau accumulation in a mouse model of tauopathy. J Exp Med. 2017;214(5):1227–1238.

37. Florian H, Wang D, Arnold SE, et al. Tilavonemab in early Alzheimer’s disease: results from a phase 2, randomized, double-blind study. Brain. 2023;146(6):2275–2284.

38. Evans HT, Benetatos J, van Roijen M, Bodea LG, Götz J. Decreased synthesis of ribosomal proteins in tauopathy revealed by non-canonical amino acid labelling. EMBO J. 2019;38(13):e101174.

39. Courade JP, Angers R, Mairet-Coello G, et al. Epitope determines efficacy of therapeutic anti-Tau antibodies in a functional assay with human Alzheimer Tau. Acta Neuropathol. 2018;136(5):729–745.

40. Albert M, Mairet-Coello G, Danis C, et al. Prevention of tau seeding and propagation by immunotherapy with a central tau epitope antibody. Brain. 2019;142(6):1736–1750.

41. Slomski A. Anti-Tau Antibody Semorinemab Fails to Slow Alzheimer Disease. JAMA. 2022;328(5):415.

42. Höglinger GU, Litvan I, Mendonca N, et al. Safety and efficacy of tilavonemab in progressive supranuclear palsy: a phase 2, randomised, placebo-controlled trial. Lancet Neurol. 2021;20(3):182–192.

43. Younes K, Sha SJ. The most valuable player or the tombstone: is tau the correct target to treat Alzheimer’s disease? Brain. 2023;146(6):2211–2213.

44. Zhou L, McInnes J, Wierda K, et al. Tau association with synaptic vesicles causes presynaptic dysfunction. Nat Commun. 2017;8:15295.

45. Combs B, Hamel C, Kanaan NM. Pathological conformations involving the amino terminus of tau occur early in Alzheimer’s disease and are differentially detected by monoclonal antibodies. Neurobiol Dis. 2016;94:18–31.

46. Stefanoska K, Volkerling A, Bertz J, et al. An N-terminal motif unique to primate tau enables differential protein-protein interactions. J Biol Chem. 2018;293(10):3710–3719.

47. Colom-Cadena M, Davies C, Sirisi S, et al. Synaptic oligomeric tau in Alzheimer’s disease - A potential culprit in the spread of tau pathology through the brain. Neuron. 2023;111(14):2170–2183.e6.

48. Hampel H, Toschi N, Babiloni C, et al. Revolution of Alzheimer Precision Neurology. Passageway of Systems Biology and Neurophysiology. J Alzheimers Dis. 2018;64(s1):S47–s105.

49. van der Ende EL, In ’t Veld S, Hanskamp I, et al. CSF proteomics in autosomal dominant Alzheimer’s disease highlights parallels with sporadic disease. Brain. 2023:awad213.

50. Chatterjee P, Pedrini S, Doecke JD, et al. Plasma Aβ42/40 ratio, p-tau181, GFAP, and NfL across the Alzheimer’s disease continuum: A cross-sectional and longitudinal study in the AIBL cohort. Alzheimers Dement. 2023;19(4):1117–1134.

51. Hirtz C, Busto GU, Bennys K, et al. Comparison of ultrasensitive and mass spectrometry quantification of blood-based amyloid biomarkers for Alzheimer’s disease diagnosis in a memory clinic cohort. Alzheimers Res Ther. 2023;15(1):34.

52. Jiang Y, Zhou X, Ip FC, et al. Large-scale plasma proteomic profiling identifies a high-performance biomarker panel for Alzheimer’s disease screening and staging. Alzheimers Dement. 2022;18(1):88–102.

53. Deutsch EW, Bandeira N, Perez-Riverol Y, et al. The ProteomeXchange consortium at 10 years: 2023 update. Nucleic Acids Res. 2023;51(D1):D1539–d1548.

54. Benjamini Y, Drai D, Elmer G, Kafkafi N, Golani I. Controlling the false discovery rate in behavior genetics research. Behav Brain Res. 2001;125(1-2):279–84.

## Supplementary references

55. Bajracharya R, Cruz E, Götz J, Nisbet RM. Ultrasound-mediated delivery of novel tau-specific monoclonal antibody enhances brain uptake but not therapeutic efficacy. J Control Release. 2022;349:634–648.

56. Liu C, Song X, Nisbet R, Götz J. Co-immunoprecipitation with Tau Isoform-specific Antibodies Reveals Distinct Protein Interactions and Highlights a Putative Role for 2N Tau in Disease. J Biol Chem. 2016;291(15):8173–88.

57. Nisbet RM, Van der Jeugd A, Leinenga G, Evans HT, Janowicz PW, Götz J. Combined effects of scanning ultrasound and a tau-specific single chain antibody in a tau transgenic mouse model. Brain. 2017;140(5):1220–1230.

58. Greenberg SG, Davies P. A preparation of Alzheimer paired helical filaments that displays distinct tau proteins by polyacrylamide gel electrophoresis. Proc Natl Acad Sci USA. 1990;87(15):5827–31.

59. Holmes BB, Furman JL, Mahan TE, et al. Proteopathic tau seeding predicts tauopathy in vivo. Proc Natl Acad Sci USA. 2014;111(41):E4376–85.

60. Polanco JC, Scicluna BJ, Hill AF, Götz J. Extracellular Vesicles Isolated from the Brains of rTg4510 Mice Seed Tau Protein Aggregation in a Threshold-dependent Manner. J Biol Chem. 2016;291(24):12445–12466.

61. Bajracharya R, Brici D, Bodea LG, Janowicz PW, Götz J, Nisbet RM. Tau antibody isotype induces differential effects following passive immunisation of tau transgenic mice. Acta Neuropathol Commun. 2021;9(1):42.

62. Ittner LM, Fath T, Ke YD, et al. Parkinsonism and impaired axonal transport in a mouse model of frontotemporal dementia. Proc Natl Acad Sci USA. 2008;105(41):15997–6002.

63. Wessel D, Flügge UI. A method for the quantitative recovery of protein in dilute solution in the presence of detergents and lipids. Anal Biochem. 1984;138(1):141–3.

64. Engholm-Keller K, Larsen MR. Improving the Phosphoproteome Coverage for Limited Sample Amounts Using TiO2-SIMAC-HILIC (TiSH) Phosphopeptide Enrichment and Fractionation. Methods Mol Biol. 2016;1355:161–77.

65. Gentleman RC, Carey VJ, Bates DM, et al. Bioconductor: open software development for computational biology and bioinformatics. Genome Biol. 2004;5(10):R80.

66. Dudoit S, Yang YH, Callow MJ, Speed TP. Statistical methods for identifying differentially expressed genes in replicated cDNA microarray experiments. Stat Sin. 2002;12(1):111–139.

67. Leek JT, Storey JD. Capturing heterogeneity in gene expression studies by surrogate variable analysis. PLoS Genet. 2007;3(9):1724–35.

68. Ritchie ME, Phipson B, Wu D, et al. limma powers differential expression analyses for RNA-sequencing and microarray studies. Nucleic Acids Res. 2015;43(7):e47.

69. Benjamini Y, Drai D, Elmer G, Kafkafi N, Golani I. Controlling the false discovery rate in behavior genetics research. Behav Brain Res. 2001;125(1-2):279–84.

70. Waardenberg AJ. Statistical Analysis of ATM-Dependent Signaling in Quantitative Mass Spectrometry Phosphoproteomics. Methods Mol Biol. 2017;1599:229–244.

71. Müller JA, Betzin J, Santos-Tejedor J, et al. A presynaptic phosphosignaling hub for lasting homeostatic plasticity. Cell Rep. 2022;39(3):110696.

72. Müller JA, Betzin J, Santos-Tejedor J, et al. A presynaptic phosphosignaling hub for lasting homeostatic plasticity. Cell Rep. 2022;39(3):110696.

73. Yu G, Wang LG, Han Y, He QY. clusterProfiler: an R package for comparing biological themes among gene clusters. Omics. 2012;16(5):284–7.

74. Zhou Y, Zhou B, Pache L, et al. Metascape provides a biologist-oriented resource for the analysis of systems-level datasets. Nat Commun. 2019;10(1):1523.

